# A GPX4 phosphorylation switch by FGFR1 guards against ferroptosis

**DOI:** 10.64898/2026.06.22.733676

**Authors:** Lintao Song, Luyao Wang, Wenliya Dong, Jie Qi, Sisi Zhi, Hongxia Lu, Yang Sun, Siyan Xu, Yushu Hou, Shuliang Tian, Jie Chen, Hao Ye, Jiaze Xi, Weinan Liang, Lei Fan, Xiaokun Li, Zhifeng Huang

## Abstract

Ferroptosis is driven by lipid peroxidation^1–3^, yet the mechanisms by which cells rapidly adjust their sensitivity to ferroptosis in response to extracellular cues remain elusive. We identify a direct phosphorylation switch controlling the activity of glutathione peroxidase 4 (GPX4), the core ferroptosis regulator^4^. The receptor tyrosine kinase (RTK) FGFR1 directly binds and phosphorylates GPX4 at Tyr180/Tyr196 in a kinase-dependent manner, requiring FGFR1 Tyr730 as a docking site. This phosphorylation enhances GPX4’s catalytic activity and suppresses ferroptosis. In cardiac ischemia/reperfusion injury, the FGFR1-GPX4 axis is suppressed, and a selective FGFR1 agonist (FGF-1^ΔNT^) reactivates it to protect against ferroptosis-mediated damage. Critically, a phosphorylation-deficient GPX4 knock-in mouse exhibits hypersensitivity to injury and is non-responsive to this agonist, demonstrating that GPX4 phosphorylation is essential. Our findings reveal a rapid mechanism for regulating ferroptosis via GPX4 tyrosine phosphorylation, directly linking RTK signaling to ferroptosis, and offering new strategies for treating ischemic-and other ferroptosis-associated diseases.

## Introduction

Ferroptosis is an iron-dependent form of regulated cell death that is driven by the accumulation of membrane lipid peroxides^3–5^. It has been implicated as a key pathological mechanism underlying a broad spectrum of human diseases, including ischemia/reperfusion (I/R) injury^6–8^, neurodegeneration^9,10^, cancer^11,12^, and acute organ damage^13^. Despite its established importance, the mechanisms by which cells dynamically modulate their sensitivity to ferroptosis in response to extracellular cues remain poorly understood.

Glutathione peroxidase 4 (GPX4) is the key mediator of the ferroptosis defense system^14,15^. As a selenoprotein, GPX4 utilizes glutathione (GSH) to reduce membrane lipid peroxides to non-toxic lipid alcohols, thereby halting the propagation of lipid peroxidation chain reactions and acting as the final gatekeeper against ferroptotic cell death^15,16^. Analyses of GPX4 regulation have primarily focused on transcriptional control (e.g., via the NRF2 pathway)^17,18^, protein stability (e.g., deubiquitinase-mediated stabilization or autophagic degradation)^19–21^, and availability of the GSH cofactor (e.g., SLC7A11-mediated cystine uptake)^4,14,22^. However, these mechanisms operate over relatively slow timescales and cannot fully account for the rapid, transient adjustments in ferroptosis sensitivity observed in response to acute extracellular signals such as growth factors or stress stimuli. To date, it is unclear whether there is a fast, reversible regulatory mechanism, such as a post-translational modification, that can directly modulate GPX4 activity.

Fibroblast growth factor receptor 1 (FGFR1) is a prototypical RTK that regulates cell proliferation, survival, and metabolism through ligand-induced dimerization and autophosphorylation^23,24^. Although FGFR1 has been implicated in the regulation of cell death^25–27^, the specific pathways it controls in this context, as well as the underlying mechanisms, are largely undefined. By systematically evaluating distinct cell death pathways, we unexpectedly discovered that FGFR1 selectively suppresses ferroptosis. Across multiple cell lines, *Fgfr1* deficiency or inhibition sensitized cells to ferroptosis inducers, but had no appreciable effect on apoptosis, necroptosis, or pyroptosis. Levels of ferroptosis were inversely correlated with FGFR1 expression levels, suggesting that FGFR1 may serve as a critical stress-responsive regulator of ferroptosis sensitivity. These findings raised a pivotal question: what are the mechanism and downstream effector by which FGFR1 regulates ferroptosis.

Here we show that GPX4 directly binds and is phosphorylated by FGFR1. This binding requires FGFR1 kinase activity, resulting in the phosphorylation of GPX4 at Tyr180 and Tyr196. Phosphorylation enhanced the catalytic activity of GPX4 by increasing its affinity for hydroperoxide substrates, thereby facilitating the clearance of membrane lipid peroxides and suppressing ferroptosis. Domain-mapping and site-directed mutagenesis analyses revealed that the FGFR1 kinase domain interacted with GPX4, with Tyr730 of FGFR1 serves as the key interface residue. In the context of cardiac I/R injury, an acute stress condition demanding rapid ferroptosis modulation, the FGFR1-GPX4 axis was suppressed, leading to GPX4 dephosphorylation and high levels of ferroptosis. To counteract this, we engineered a selective FGFR1 agonist, FGF-1^ΔNT^, which activated this axis, suppressed ferroptosis, and attenuated myocardial injury. Crucially, a phosphorylation-deficient *Gpx4* knock-in mouse (*Gpx4*-Y180/196F) exhibited heightened sensitivity to ferroptosis-induced damage, and FGF-1^ΔNT^ administration to this mouse failed to provide a protective effect. Collectively, these findings establish a direct mechanistic link between RTK signaling and ferroptosis execution through the phosphorylation of GPX4. This represents a rapid-response regulatory mechanism for ferroptosis and offers a potential strategy for treating ischemic heart disease and other ferroptosis-associated pathologies.

## Results

### FGFR1 selectively suppresses ferroptosis

Recent studies have implicated FGFR1 in the regulation of cell death^25–27^. However, the specific types of cell death it modulates, and the underlying mechanisms remain poorly defined. To systematically assess the role of FGFR1 in different cell death pathways, we evaluated its function in mouse embryonic fibroblasts (MEFs) subjected to apoptosis, necroptosis, pyroptosis or ferroptosis. *Fgfr1* deficiency markedly exacerbated lipid peroxidation and ferroptosis induced by Erastin, RSL3, or cystine depletion (**Fig. 1a, b** and **Extended Data Fig. 1a, b**). By contrast, *Fgfr1* deficiency had minimal effects on apoptosis, necroptosis or pyroptosis induced by a range of stimuli (**Extended Data Fig. 1c-f**). Consistent with this result, *FGFR1* deletion increased the sensitivity of HT1080 cells to Erastin and RSL3 (**Fig. 1c-e** and **Extended Data Fig. 1g-k**), and reintroduction of FGFR1 into *FGFR1* knockout (*FGFR1*^−/−^) HT1080 cells effectively attenuated lipid peroxidation and ferroptosis in response to these stimuli (**Fig. 1f-h**). Furthermore, the ferroptosis inhibitors ferrostatin-1 (Fer-1) and the iron chelator deferoxamine (DFO), but not inhibitors of necroptosis (necrostatin-1, Nec) or apoptosis (Z-VAD-FMK, ZVF), abolished the cell death and lipid peroxidation induced by RSL3 or Erastin in *Fgfr1*-deficient cells (**Fig. 1i, j**).

**Fig. 1.**
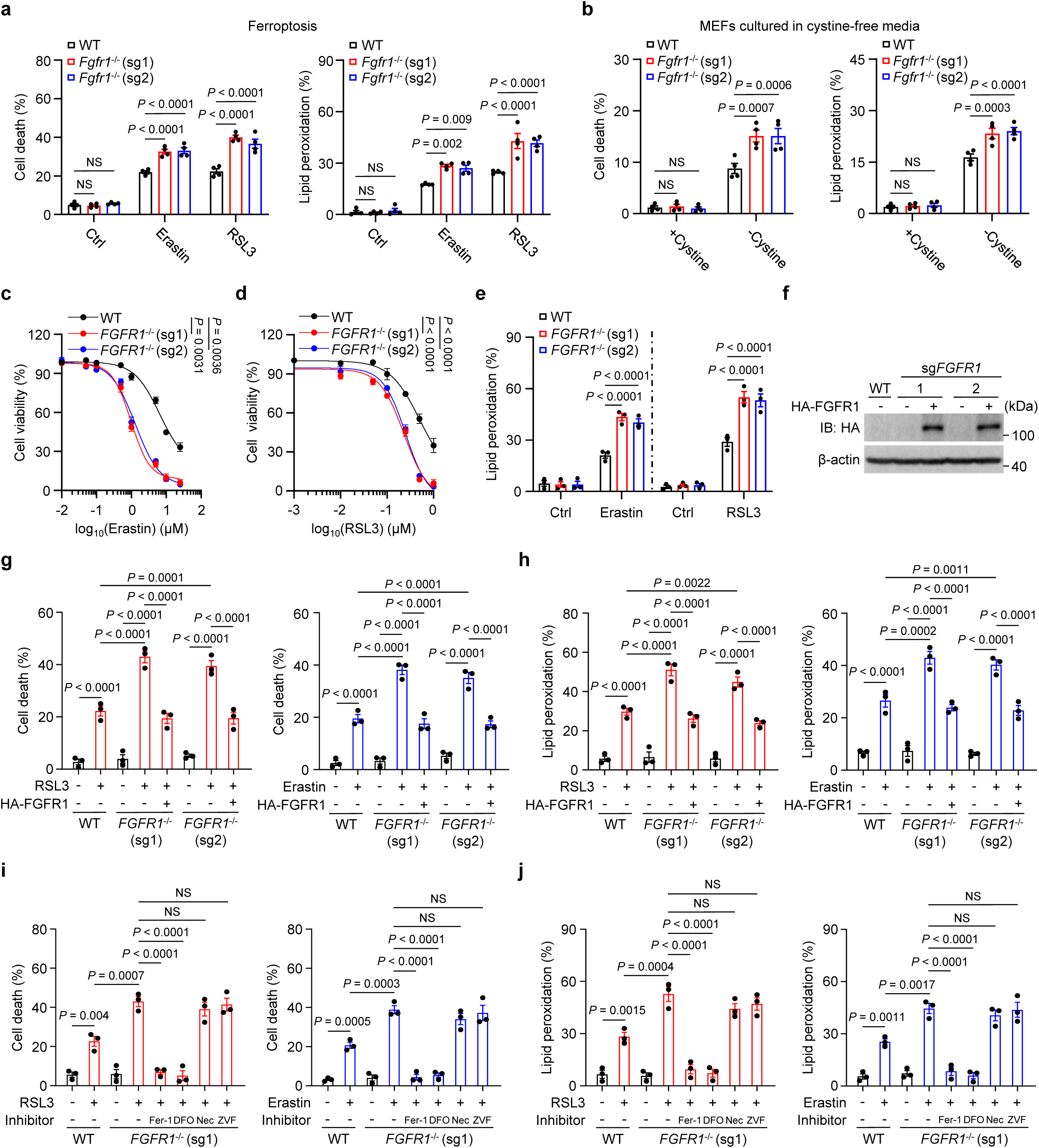
FGFR1 is a potent ferroptosis suppressor. **a,** Cell death (left) and lipid peroxidation (right) were performed with wild-type (WT) and *Fgfr1-*deficient (*Fgfr1*^−/−^, sg1 or sg2) MEFs treated with 10 μM Erastin or 0.5 μM RSL3. **b,** Cell death (left) and lipid peroxidation (right) in WT and *Fgfr1*^−/−^MEFs treated with or without 0.1 mM cystine in cystine-free media for 9 h. **c,d,** Cell viability in WT and *FGFR1* deficient (*FGFR1*^−/−^, sg1 or sg2) HT1080 cells treated with or without Erastin (**c**) or RSL3 (**d**). **e,** Lipid-peroxidation measurements in the indicated HT1080 cells treated with 10 μM Erastin or 0.5 μM RSL3. **f,** Immunoblot analysis of FGFR1 levels in the indicated HT1080 cells. *FGFR1*^−/−^ (sg1 and sg2) HT1080 cells were infected with empty vector or HA-tagged FGFR1. **g,h,** Cell death (**g**) and lipid peroxidation (**h**) in WT and *FGFR1*^−/−^ (sg1 or sg2) HT1080 cells reconstituted with empty vector or HA-tagged FGFR1 and treated with 0.5 μM RSL3 or 10 μM Erastin. **i,j,** Cell death (**i**) and lipid peroxidation (**j**) measurements in WT and *FGFR1*^−/−^ (sg1) HT1080 cells subjected to the indicated treatments. RSL3, 0.5 μM; Erastin, 10 μM; Fer-1 and DFO, 5 μM; Nec, 2 μM necrostatin-1; ZVF, 10 μM Z-VAD-FMK. *n* = 3-4 biologically independent experiments in (**a-e, g-j**). Data are mean ± s.e.m.; ordinary two-way ANOVA, followed by Sidak in (**a, b, e**); two-way ANOVA (repeated measure), followed by Tukey in (**c, d**); ordinary one-way ANOVA, followed by Tukey in (**g-j**); \*\**p* < 0.01, \*\*\**p* < 0.001, \*\*\*\**p* < 0.0001, NS, not significant; Gel source data blots can be found in Supplementary Fig. 1.

To assess whether FGFR1-mediated ferroptosis regulation is a generalizable phenomenon, we compared basal levels of FGFR1 expression to ferroptosis sensitivity across a panel of cell lines (**Extended Data Fig. 2a**). Cells with low FGFR1 expression exhibited higher sensitivity to ferroptosis inducers than those with high FGFR1 expression (**Extended Data Fig. 2b, c**). Knockdown of *FGFR1* in FGFR1-high cell lines (U251, MDA-MB-231 and NCI-H1299) enhanced Erastin-and RSL3-induced lipid peroxidation and ferroptosis (**Extended Data Fig. 2d-f**). Conversely, overexpression of FGFR1 in FGFR1-low cell lines (HeLa, HepG2, Huh7 and NCI-H1975) dramatically attenuated ferroptosis and lipid peroxidation (**Extended Data Fig. 2g-j**). Collectively, these data indicate that FGFR1 specifically modulates cellular sensitivity to ferroptosis, establishing it as a critical suppressor of this cell death pathway.

### FGFR1 kinase activity is essential for ferroptosis resistance

Given that FGFR1 functions as an RTK^24^, we therefore investigated whether its kinase activity is required for ferroptosis suppression. Treatment of HT1080 and MEF cells with Erastin or RSL3 inhibited FGFR1 kinase activity in a concentration- and time-dependent manner, as evidenced by reduced autophosphorylation of Tyr653 and Tyr654 within the activation loop of the FGFR1 kinase domain^28^ (**Extended Data Fig. 3a-f**). To validate these findings, we treated cells with specific FGFR1 inhibitors, namely PD166866 and PD173074. Pharmacological inhibition of FGFR1 significantly enhanced Erastin-or RSL3-induced lipid peroxidation and ferroptosis; notably, this effect was observed exclusively in FGFR1-high cells (**Extended Data Fig. 4a-f**). In FGFR1-high cells, co-treatment with ferroptosis inducers and an FGFR1 inhibitor resulted in levels of cell death and lipid peroxidation comparable to those seen in FGFR1-low cells; this effect was abrogated by the ferroptosis inhibitor Fer-1 (**Extended Data Fig. 4c-f**). More importantly, in *FGFR1*-deficient HT1080 cells, re-introduction of wild-type FGFR1 (WT-FGFR1), but not a kinase-dead mutant (KD-FGFR1, Y653/654F), restored resistance to Erastin-and RSL3-induced ferroptosis and lipid peroxidation (**Fig. 2a**). These data demonstrate that the anti-ferroptotic function of FGFR1 strictly depends on its kinase activity.

**Fig. 2.**
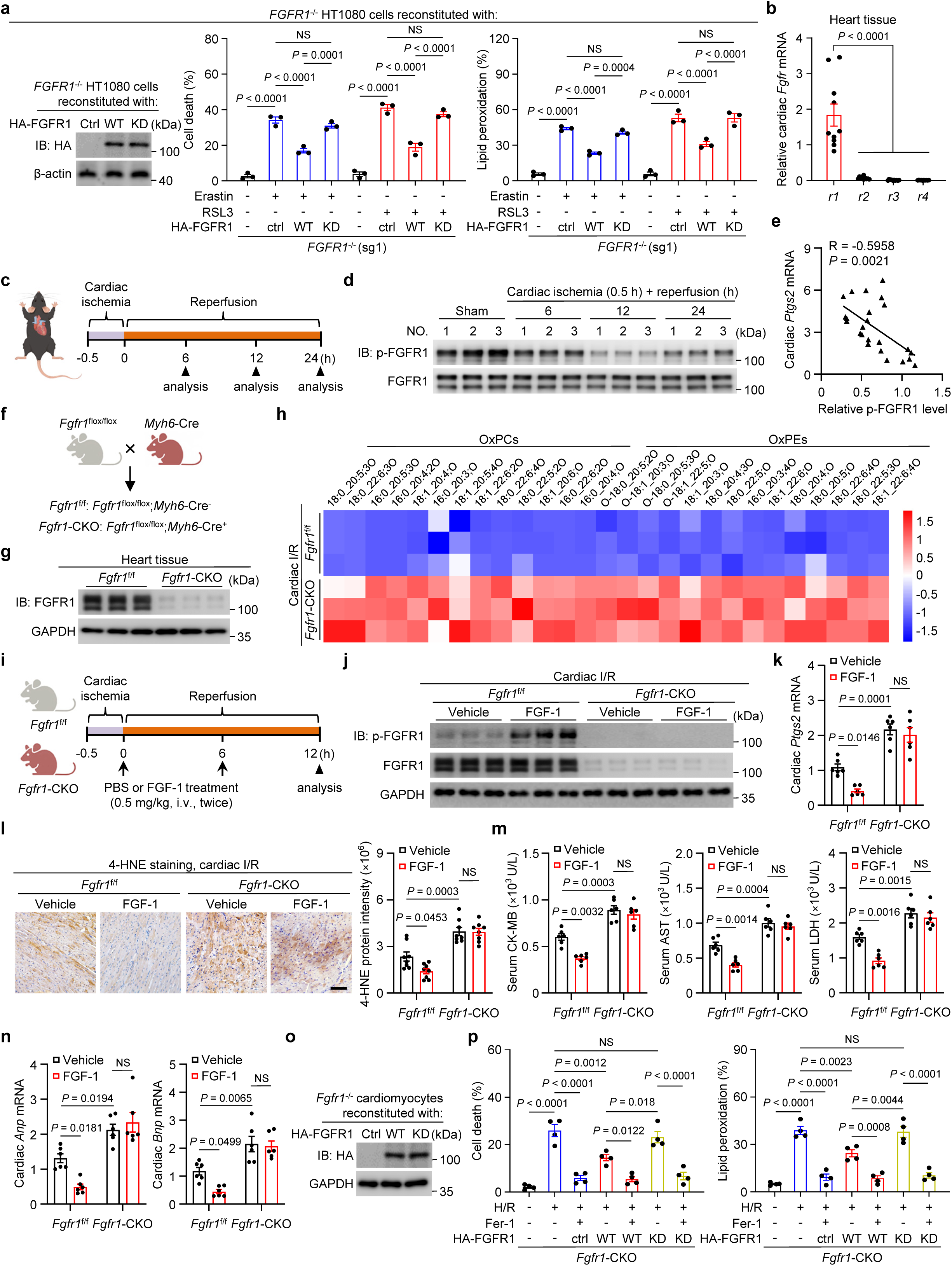
FGFR1 kinase activity is essential for ferroptosis triggered by cardiac ischemia-reperfusion (I/R) injury. **a,** Cell death and lipid peroxidation were performed with *FGFR1*^−/−^ HT1080 cells reconstituted with empty vector (Ctrl) or HA-tagged WT-FGFR1, or KD-FGFR1 (a kinase-dead form of FGFR1, FGFR1-Y653/654F, Tyr653 and Tyr654 are important for the catalytic activity of activated FGFR1) subjected to 10 μM Erastin or 0.5 μM RSL3 treatments. **b,** Relative mRNA levels of *Fgfr1-4* in cardiac tissues of C57BL/6J mice analyzed by qRT-PCR. Relative mRNA levels were normalized to *Gapdh*. *n* = 10 mice/group. **c,** Schematic representation of cardiac I/R in male C57BL/6J mice. The mice were subjected to 0.5 h of ischemia followed by 6, 12, or 24 h of reperfusion, with a sham group serving as control. **d,** Immunoblot analysis of phosphorylated FGFR1 (Tyr653/654, p-FGFR1) levels in total cardiac lysates from C57BL/6J mice collected at the indicated time after cardiac I/R treatment. **e,** Correlation between cardiac *Ptgs2* mRNA levels and relative p-FGFR1 in cardiac tissues from the indicated mice. Ratios of p-FGFR1 to total FGFR1 were calculated. Relative mRNA levels were normalized to *Gapdh*. *n* = 24 mice. **f,** Schematic of cardiac-specific *Fgfr1* deficiency (*Fgfr1*-CKO) mice. **g,** Immunoblot analysis confirming successful cardiac-specific *Fgfr1* knockout in mice. **h,** Heat map showing relative amounts of oxidized phospholipids in the heart tissues collected 12 h after cardiac I/R. *n* = 3 mice/group. **i,** Schematic of the treatment regimen with FGF-1 in cardiac I/R-treated *Fgfr1*^f/f^ and cardiac-specific *Fgfr1*-deficient (*Fgfr1*-CKO) mice. Indicated mice were subjected to cardiac I/R, followed by intravenous (i.v.) administration of FGF-1 (0.5 mg/kg) at the onset of reperfusion and 6 h post-reperfusion. **j,** Immunoblot analysis of total cardiac samples from the indicated mice. **k,** Relative *Ptgs2* mRNA levels in the indicated mice. *n* = 6 mice/group. **l,** Representative images and quantification of 4-HNE– stained cardiac sections from the indicated mice. Scale bar, 100 μm. *n* = 8 mice/group. **m,** Serum creatine kinase-MB isoenzyme (CK-MB), aspartate transaminase (AST), and lactate dehydrogenase (LDH) levels in cardiac I/R-treated *Fgfr1*^f/f^ and *Fgfr1*-CKO mice upon FGF-1 treatment. *n* = 6 mice/group. **n,** Relative mRNA levels of *Anp* and *Bnp* in the indicated mice. *n* = 6 mice/group. **o,** Immunoblot showing FGFR1 levels in *Fgfr1*-CKO cardiomyocytes infected with empty vector (ctrl), HA-WT-FGFR1 or KD-FGFR1. **p,** Cell death (left) and lipid peroxidation (right) were performed with *Fgfr1*-CKO cardiomyocytes reconstituted with empty vector (ctrl), HA-tagged WT-FGFR1 or KD-FGFR1 subjected to hypoxia/reoxygenation (H/R) challenge in the presence or absence of ferroptosis inhibitors Fer-1. *n* = 3-4 biologically independent experiments in (middle and right graphs of **a, p**). Data are mean ± s.e.m.; ordinary one-way ANOVA, followed by Tukey in (middle and right graphs of **a, b, p**); Pearson’s correlation in (**e**); ordinary two-way ANOVA, followed by Tukey in (**k, m, n,** right graph of **l**); \**p* < 0.05, \*\**p* < 0.01, \*\*\**p* < 0.001, \*\*\*\**p* < 0.0001, NS, not significant; Gel source data blots can be found in Supplementary Fig. 1.

Ferroptosis is a major driver of cell death in myocardial ischemia/reperfusion (I/R) injury^8,29^, and FGFR1 is the predominant FGFR subtype expressed in heart tissue^30,31^ (**Fig. 2b**). To explore the pathological relevance of FGFR1-mediated ferroptosis suppression in this context, we employed a mouse model of cardiac I/R injury in C57BL/6J mice (**Fig. 2c**). I/R injury resulted in elevated levels of cardiac enzymes (CK-MB, AST and LDH) in serum (**Extended Data Fig. 5a-c**). In heart tissue, this was accompanied by increased expression of cardiac injury markers (*Anp* and *Bnp*), along with time-dependent upregulation of ferroptosis markers (*Ptgs2* mRNA and 4-HNE protein) (**Extended Data Fig. 5d-f**). Notably, tyrosine phosphorylation of FGFR1 (p-Tyr653/654) in heart tissue was significantly suppressed after I/R, reaching its lowest level after 12 h of reperfusion (**Fig. 2d**), and p-FGFR1 signal intensity was inversely correlated with ferroptosis maker expression (**Fig. 2e** and **Extended Data Fig. 5g**). Consistent with these *in vivo* observations, hypoxia/reoxygenation (H/R) treatment of primary mouse cardiomyocytes also led to reduced p-FGFR1 levels (**Extended Data Fig. 5h, i**).

To further explore the causal role of FGFR1 in cardiac I/R injury, we generated cardiomyocyte-specific *Fgfr1* knockout (*Fgfr1*-CKO) mice (**Fig. 2f, g** and **Extended Data Fig. 5j**). Compared with *Fgfr1*^f/f^ controls, *Fgfr1*-CKO mice exhibited exacerbated cardiac injury after I/R. Lipidomic profiling of heart tissues collected 12 h after I/R revealed a global increase in oxidized phospholipids in *Fgfr1*-CKO mice (**Fig. 2h**). In line with this, *Fgfr1*-CKO mice showed increased lipid peroxidation and ferroptosis markers (**Fig. 2i-l** and **Extended Data Fig. 5k, l**), significantly higher levels of serum CK-MB, AST and LDH, elevated *Anp* and *Bnp* mRNA expression (**Fig. 2m, n**). By contrast, *Fgfr1* deficiency had minimal effects on apoptosis, necroptosis, or pyroptosis induced by cardiac I/R (**Extended Data Fig. 5m**). Administration of FGF-1, a natural ligand of FGFR1, restored FGFR1 phosphorylation in heart tissue of *Fgfr1*^f/f^ mice subjected to I/R injury and markedly ameliorated I/R-induced ferroptosis and cardiac injury (**Fig. 2i-n** and **Extended Data Fig. 5k, l**). Crucially, this protective effect of FGF-1 was completely abolished in *Fgfr1*-CKO mice (**Fig. 2j-n** and **Extended Data Fig. 5k, l**). Consistent with this result, re-introduction of WT-FGFR1, but not KD-FGFR1, into *Fgfr1*-deficient primary cardiomyocytes restored resistance to H/R-induced ferroptosis (**Fig. 2o, p**). Notably, treatment with the ferroptosis inhibitors Fer-1 or liproxstatin-1 (Lip-1) restored FGFR1 phosphorylation in both H/R-treated cardiomyocytes and I/R-challenged heart tissue (**Extended Data Fig. 5n-p**). Together, these findings support an essential role for FGFR1 kinase activity is critical in suppressing ferroptosis during myocardial I/R injury.

### FGFR1 regulates ferroptosis through direct interactions with GPX4

Canonical FGFR signaling involves ligand-induced dimerization, autophosphorylation, and the recruitment of downstream signaling proteins including the effector PLCγ and the adaptor protein FRS2^32,33^ (**Fig. 3a**). To test whether these canonical pathways mediate the anti-ferroptotic effect of FGFR1, we used CRISPR/Cas9 to delete *Plcγ* or *Fr*s2 in MEFs and assessed their sensitivity to ferroptosis and lipid peroxidation (**Fig. 3b** and **Extended Data Fig. 6a, b**). Contrary to expectations, loss of either *Plcγ* or *Frs2* did not phenocopy *Fgfr1* deficiency; neither *Plcγ* nor *Frs2* ablation enhanced Erastin- or RSL3-induced ferroptosis or lipid peroxidation accumulation (**Fig. 3c** and **Extended Data Fig. 6c, d**). In addition, neither *FGFR1* knockout nor FGFR1 inhibition affected cellular divalent iron levels, excluding the possibility that FGFR1 acts by altering iron availability (**Extended Data Fig. 6e, f**). These findings suggest that FGFR1 regulates ferroptosis through an alternative, non-canonical pathway involving an unidentified substrate.

**Fig. 3.**
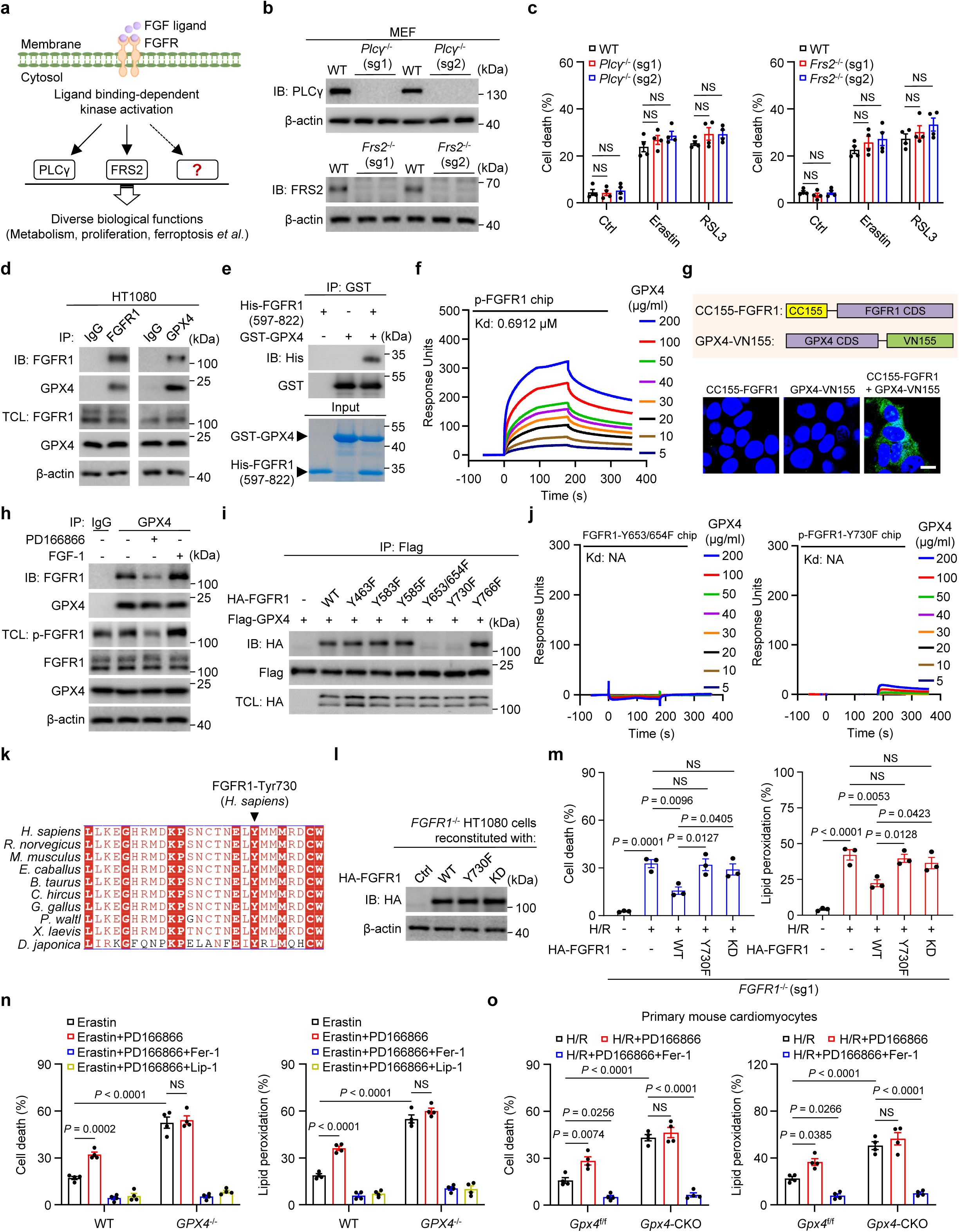
FGFR1 regulates ferroptosis by directly interacting with GPX4. **a,** Simplified overview of FGFR1 signaling pathways. FGF ligand binding induces FGFR1 dimerization and autophosphorylation. Phosphorylated tyrosine residues serve as docking sites for PLCγ and FRS2, leading to cellular proliferation, metabolism, and ferroptosis. **b,** Immunoblot analysis of PLCγ and FRS2 protein levels in CRISPR control (WT) and knockout (*Plcγ*^−/−^ or *Frs2*^−/−^) MEFs. **c,** Cell death in WT and *Plcγ*-deficient (left) or *Frs2*-deficient (right) MEFs treated with 10 μM Erastin or 0.5 μM RSL3. **d,** Interaction between endogenous FGFR1 and GPX4 in HT1080 cells. Immunoprecipitation (IP) of FGFR1 or GPX4 followed by immunoblotting of co-precipitated proteins. TCL, total cell lysate. **e,** GST pull-down assay using bacterially purified His-FGFR1 (aa 597-822) and GST-GPX4. **f,** Surface plasmon resonance (SPR) analysis of GPX4 binding to phosphorylated FGFR1 tyrosine kinase domain. Equilibrium dissociation constants (Kd) were derived from saturation binding curves. **g,** Bimolecular fluorescence complementation (BiFC) assay showing FGFR1 and GPX4 colocalization in HEK293T cells. CDS, coding sequence. Scale bars, 10 μm. **h,** Effect of FGFR1 inhibition or activation on FGFR1-GPX4 interaction. HT1080 cells were treated with PD166866 (FGFR1 inhibitor) for 4 h, or FGF-1 for 0.5 h. **i,** Identification of FGFR1 Tyr730 as critical for GPX4 binding. HA-tagged FGFR1 mutants (tyrosine to phenylalanine) were co-expressed with Flag-GPX4 in HEK293T cells. Seven tyrosine residues in the cytoplasmic tail of FGFR1 can be phosphorylated: Tyr463, 583, 585, 653, 654, 730, and 766. **j,** The FGFR1-Y653/654F (KD, left) and FGFR1-Y730F (right) mutants fail to interact with GPX4 by SPR. **k,** Sequence alignment of the residues flanking Tyr730 of FGFR1 across different species. Arrow heads point to Tyr730 corresponding to human FGFR1. **l,** Immunoblot showing the expression of FGFR1 in *FGFR1*^−/−^ HT1080 cells infected with empty vector (ctrl), and HA-tagged WT-FGFR1, Y730F-FGFR1 or KD-FGFR1. **m,** Cell death and lipid peroxidation were performed in *FGFR1*^−/−^ HT1080 cells reconstituted with empty vector, and HA-tagged WT-FGFR1, Y730F-FGFR1 or KD-FGFR1 subjected to H/R treatments. **n,o,** Measurement of cell death and lipid peroxidation in WT and *GPX4*^−/−^HT1080 cells treated with Erastin (**n**), or *Gpx4*^f/f^ and *Gpx4*-CKO cardiomyocytes upon H/R (**o**), with or without the indicated inhibitors. *n* = 3-4 biologically independent experiments in (**c, m-o**). Data are mean ± s.e.m.; ordinary two-way ANOVA, followed by Sidak in (**c**); ordinary one-way ANOVA, followed by Tukey in (**m**); ordinary two-way ANOVA, followed by Tukey in (**n, o**); \**p* < 0.05, \*\**p* < 0.01, \*\*\**p* < 0.001, \*\*\*\**p* < 0.0001, NS, not significant; Gel source data blots can be found in Supplementary Fig. 1.

To identify this substrate, we Flag-tagged endogenous FGFR1 in cardiomyocytes (**Extended Data Fig. 7a**) and performed immunoprecipitation followed by mass spectrometry. This identified GPX4 as a potential interacting partner (**Extended Data Fig. 7b**). A direct physical association between FGFR1 and GPX4 was confirmed by co-immunoprecipitation (in HT1080 and HEK293T cells), GST pull-down assays and surface plasmon resonance (SPR) experiment with purified proteins, bimolecular fluorescence complementation (BiFC), and proximity ligation assay (PLA) experiments (**Fig. 3d-g** and **Extended Data Fig. 7c, d**). Importantly, genetic ablation of *Plcγ* and *Frs2* did not affect this interaction (**Extended Data Fig. 7e, f**). PLA and co-immunofluorescence further revealed that FGFR1 and GPX4 co-localize, at least in part, at the plasma membrane (**Extended Data Fig. 7d, g**). Notably, this interaction strictly depended on FGFR1 kinase activity, as FGFR1 pharmacological inhibition (with PD166866 or PD173074) or expression of a kinase-dead FGFR1 mutant (KD-FGFR1), abolished the association (**Fig. 3h** and **Extended Data Fig. 7h, i**), whereas FGF-1 stimulation markedly enhanced the co-precipitation of GPX4 with FGFR1 (**Fig. 3h** and **Extended Data Fig. 7d**). Domain mapping also revealed that the association was mediated exclusively by the kinase domain of FGFR1 (**Extended Data Fig. 7j, k**). Collectively, these data provide strong evidence that FGFR1 directly binds to GPX4 in a kinase-activity-dependent manner.

Previous structural and functional studies have identified seven FGFR1 autophosphorylation sites (Tyr463, Tyr583, Tyr585, Tyr653, Tyr654, Tyr730, and Tyr766) that are required for recruiting downstream effectors^23,32–34^. Using site-directed mutagenesis, we found that a kinase-dead FGFR1 mutant (KD-FGFR1, Y653F/Y654F) failed to bind GPX4 (**Fig. 3i**). In addition, mutation of the Tyr730 site (Y730F) also abolished this interaction, whereas mutation of the other four tyrosine residues (Y463F, Y583F, Y585F, or Y766F) did not affect binding (**Fig. 3i**). This finding was independently validated by SPR experiments (**Fig. 3j**). Notably, single mutation of Tyr730 did not alter FGF-1-induced activation of FGFR1/ERK1/2 signaling (**Extended Data Fig. 8a**), consistent with a previous report^35^, suggesting that Y730F selectively impairs GPX4 binding rather than overall receptor function. Sequence alignment showed that Tyr730 of FGFR1 is highly conserved across species (**Fig. 3k**). Reconstituting *FGFR1*^−/−^ HT1080 cells with either KD-FGFR1 or the Y730F mutant failed to restore resistance to H/R-or Erastin-induced ferroptosis or to suppress lipid peroxidation (**Fig. 3l, m** and **Extended Data Fig. 8b**). These results demonstrate that both FGFR1 catalytic activity and the integrity of the Tyr730 residue are essential for GPX4 interaction and subsequent ferroptosis suppression.

To establish that the anti-ferroptotic effect of FGFR1 is mediated by GPX4, we treated control and *Gpx4*-deficient MEFs or HT1080 cells with the FGFR1 inhibitor PD166866. As expected, *Gpx4* knockdown (shRNA) or knockout (sgRNA) markedly sensitized cells to Erastin-induced ferroptosis and lipid peroxidation (**Fig. 3n** and **Extended Data Fig. 8c-f**). Notably, adding PD166866 to these cells failed to further enhance ferroptosis (**Fig. 3n** and **Extended Data Fig. 8d**), indicating that FGFR1 inhibition does not affect ferroptosis when GPX4 is absent. This epistatic relationship was validated in primary mouse cardiomyocytes subjected to H/R; FGFR1 inhibition failed to exacerbate ferroptosis in *Gpx4* conditional knockout (*Gpx4*-CKO) cardiomyocytes (**Fig. 3o** and **Extended Data Fig. 8g**). Thus, GPX4 is an essential downstream effector through which FGFR1 suppresses ferroptosis.

### Tyrosine phosphorylation of GPX4 by FGFR1 is essential for GPX4 activity and ferroptosis suppression

We next asked whether FGFR1 directly phosphorylates GPX4 on tyrosine residues. Co-expression of HA-FGFR1 with Flag-GPX4 resulted in robust tyrosine phosphorylation of GPX4, which was completely abolished by calf intestinal alkaline phosphatase (CIP) (**Fig. 4a**). Unlike wild-type FGFR1, a kinase-dead mutant (KD-FGFR1) failed to phosphorylate GPX4 *in vitro* (**Fig. 4b**); this phosphorylation was also blocked by the FGFR1 inhibitor PD166866 (**Fig. 4c**). Conversely, FGF-1 stimulation markedly enhanced the tyrosine phosphorylation of GPX4, accompanied by increased phosphorylation of the FGFR1 activation sites (Tyr653/654) (**Fig. 4c** and **Extended Data Fig. 9a**). Notably, in *FGFR1*-deficient MEF or HT1080 cells, FGF-1 treatment failed to induce GPX4 tyrosine phosphorylation (**Fig. 4c** and **Extended Data Fig. 9a**). Given the functional redundancy within the FGFR family, we also examined other widely expressed FGFR members and found that GPX4 phosphorylation was specific to FGFR1 (**Extended Data Fig. 9b**). Together with the direct FGFR1-GPX4 interaction established above, these results identify GPX4 as a tyrosine-phosphorylated substrate of FGFR1 and indicate that this phosphorylation depends on FGFR1 kinase activity.

**Fig. 4.**
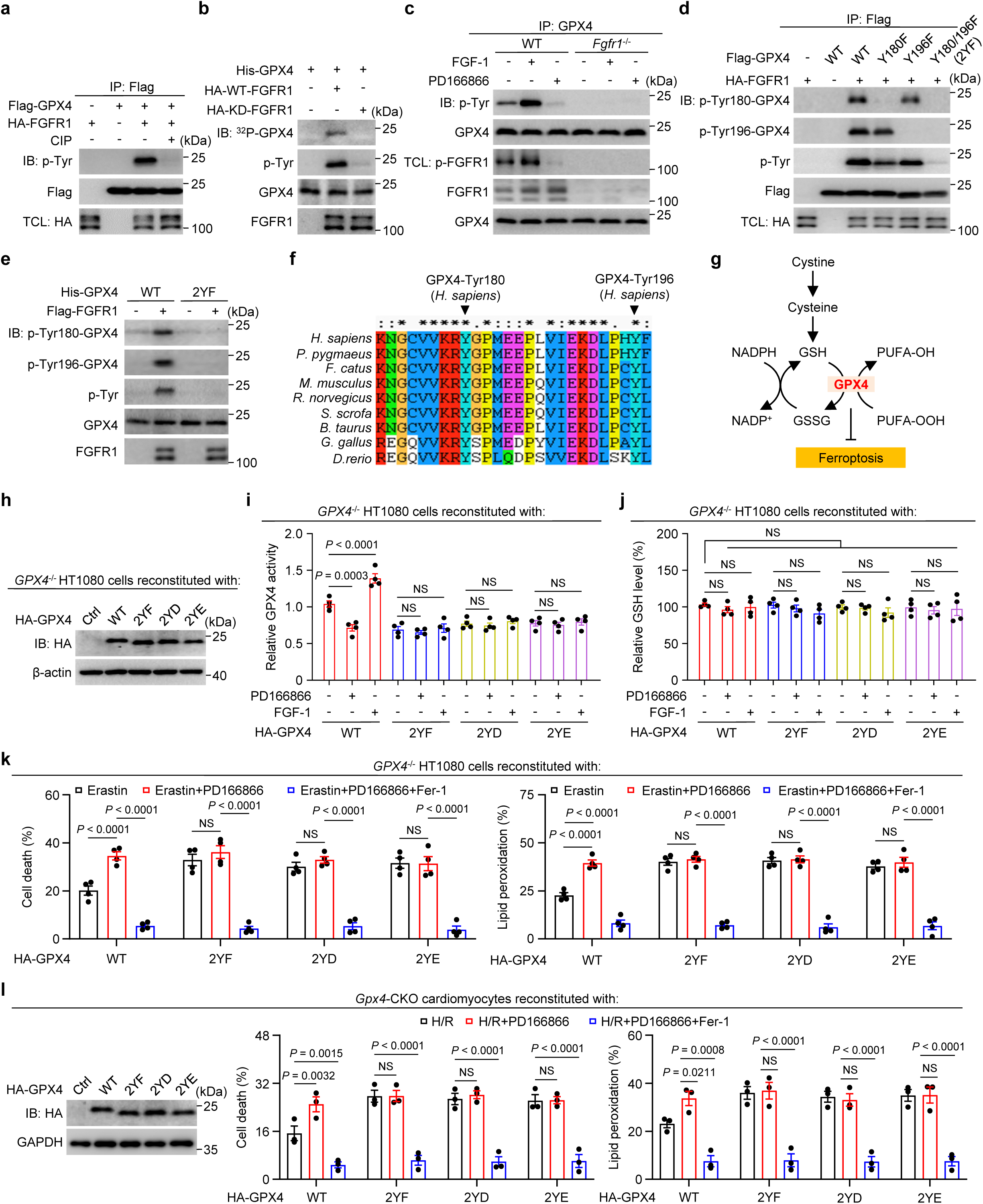
FGFR1-mediated tyrosine phosphorylation of GPX4 is required for its activity and suppresses ferroptosis. **a,** HEK293T cells were co-transfected with Flag-GPX4 and HA-FGFR1, followed by IP of Flag-GPX4 and immunoblotting with a pan anti-phospho-tyrosine (p-Tyr) antibody to detect GPX4 phosphorylation. CIP, calf-intestinal alkaline phosphatase. **b,** *In vitro* kinase assay confirming direct phosphorylation of GPX4 by FGFR1. HA-tagged WT-and KD-FGFR1 were immunoprecipitated from transfected HEK293T cells and incubated with bacterially expressed His-GPX4 in a kinase assay buffer, followed by immunoblotting. **c,** Endogenous GPX4 tyrosine phosphorylation levels after treatment with FGFR1 inhibitor PD166866 or the FGFR1 ligand FGF-1. WT or *Fgfr1*^−/−^ MEFs were treated with PD166866 for 4 h, or FGF-1 for 0.5 h. Endogenous GPX4 was immunoprecipitated and analyzed by immunoblotting. **d,** Identification of GPX4 phosphorylation sites targeted by FGFR1. Flag-WT-GPX4 or tyrosine-to-phenylalanine (YF) mutants were co-expressed with or without HA-FGFR1 in HEK293T, immunoprecipitated using an anti-Flag antibody, and immunoblotted as indicated. **e,** The Y180/196F-GPX4 mutant fails to be phosphorylated by FGFR1 *in vitro*. Flag-tagged FGFR1 proteins were isolated from HEK293T cells using anti-Flag antibody. Bacterially expressed His-tagged WT-GPX4 or its 2YF mutant was then incubated with the isolated Flag-FGFR1 in a kinase assay buffer, followed by immunoblotting. **f,** Sequence alignment of residues flanking Tyr180 and Tyr196 across different species. Arrowheads indicate Tyr180 and Tyr196 in human GPX4. **g,** Schematic diagram of GPX4 functioning as a glutathione-dependent suppressor of ferroptosis. **h,** Immunoblot analysis of proteins of *GPX4*^−/−^ HT1080 cells infected with vector (ctrl), WT-GPX4 or its 2YF, 2YD and 2YE mutants. **i,j,** GPX4 activity (**i**) or relative glutathione (GSH) levels (**j**) in *GPX4*^−/−^ HT1080 cells re-expressed HA-tagged WT-GPX4, the phosphorylation-defective 2YF, or the phosphomimetic (2YD and 2YE) mutants of GPX4. Cells were incubated with or without FGFR1 inhibitor PD166866 or the ligand FGF-1. GPX4 was immunoprecipitated using an anti-HA antibody, and GPX4 activity was measured with tert-butyl hydroperoxide (TBHP) as the substrate. **k,** Cell death (left) and lipid peroxidation (right) in *GPX4*^−/−^ HT1080 cells reconstituted with vector (ctrl), WT-GPX4, or indicated mutants and subjected to the indicated treatments. **l,** Immunoblot analysis (left) and quantification of cell death (middle) and lipid peroxidation (right) in *Gpx4*-CKO cardiomyocytes reconstituted with WT-GPX4 or its mutants and subjected to the indicated treatments. *n* = 3-4 biologically independent experiments in (**i-l**). Data are mean ± s.e.m.; ordinary one-way ANOVA, followed by Tukey in (**i, j**); ordinary two-way ANOVA, followed by Sidak in (**k, l**); \**p* < 0.05, \*\**p* < 0.01, \*\*\**p* < 0.001, \*\*\*\**p* < 0.0001, NS, not significant; Gel source data blots can be found in Supplementary Fig. 1.

To map the specific tyrosine residues targeted by FGFR1, we used the GPS 6.0 program to predict potential phosphorylation sites on GPX4. This analysis yielded six candidate tyrosine residues (**Extended Data Fig. 9c**, **d**). Site-directed mutagenesis revealed that the combined mutation of GPX4 Tyr180 and Tyr196 (2YF) dramatically reduced FGFR1-catalyzed GPX4 tyrosine phosphorylation (**Fig. 4d, e**). Sequence alignment showed that Tyr180 and Tyr196 are highly conserved across species (**Fig. 4f**). To validate these sites, we generated phospho-specific polyclonal antibodies that recognize GPX4 phosphorylated at Tyr180 or Tyr196 (**Fig. 4d, e** and **Extended Data Fig. 9e**, **f**). Using these antibodies, we confirmed that both residues are indeed phosphorylated in an FGFR1-dependent manner in cultured cells and *in vitro* (**Fig. 4d, e** and **Extended Data Fig. 9g**). Furthermore, in *GPX4*^−/−^ HT1080 cells expressing the phosphorylation-deficient 2YF-GPX4 mutant, GPX4 p-Tyr signals were almost undetectable following FGF-1 stimulation (**Extended Data Fig. 9g**). These data define Tyr180 and Tyr196 as the principal FGFR1-dependent tyrosine phosphorylation sites on GPX4.

To determine whether this phosphorylation regulates GPX4 catalytic activity, we measured GPX4 activity by monitoring the rate of NADPH oxidation (**Fig. 4g**), with *tert*-butyl hydroperoxide (TBHP, a general peroxide substrate) or phosphatidylcholine hydroperoxide (PCOOH, a physiological GPX4 substrate). Both genetic and pharmacological inhibition of FGFR1 significantly reduced GPX4 enzymatic activity in HT1080 cells under basal conditions and in the presence of the ferroptosis inducer RSL3 (**Extended Data Fig. 9h, i**). Reconstitution of *FGFR1*^−/−^ HT1080 cells with HA-FGFR1 fully restored GPX4 activity (**Extended Data Fig. 9i**), indicating that GPX4 function depends on FGFR1 signaling. Importantly, the 2YF-GPX4 mutant exhibited lower enzymatic activity than wild-type GPX4 in HEK293T and HT1080 cells (**Fig. 4h, i** and **Extended Data Fig. 9j, k**). Moreover, FGF-1 stimulation or FGFR1 overexpression increased the activity of wild-type GPX4 but failed to enhance that of the 2YF mutant (**Fig. 4i** and **Extended Data Fig. 9j, k**). These results suggest that FGFR1-mediated phosphorylation at Tyr180/Tyr196 is required for full GPX4 catalytic activity. Interestingly, phosphomimetic GPX4 mutants (2YD and 2YE) showed activity levels comparable to those of the non-phosphorylatable 2YF mutant (**Fig. 4i** and **Extended Data Fig. 9k**), indicating that glutamic acid or aspartic acid substitutions cannot functionally recapitulate FGFR1-mediated tyrosine phosphorylation of GPX4. Kinetic analysis further revealed that FGFR1-mediated phosphorylation enhances GPX4 activity by increasing its binding affinity for the TBHP substrate (**Extended Data Fig. 9l**). Consistent with previous reports^14,36^, GSH levels remained largely unchanged after RSL3 treatment or *GPX4* deficiency, and neither FGFR1 activation/inhibition nor mutation of GPX4 altered GSH levels (**Fig. 4j** and **Extended Data Fig. 9m, n**), suggesting that FGFR1 regulates GPX4 activity via Tyr180/Tyr196 phosphorylation rather than through GSH availability.

We then examined whether this phosphorylation-dependent activation of GPX4 influences ferroptosis sensitivity. Compared with cells reconstituted with wild-type GPX4, *GPX4*^−/−^ HT1080 cells expressing the 2YF, 2YD, or 2YE mutants exhibited increased lipid peroxidation and ferroptosis upon Erastin treatment (**Fig. 4k**). In *GPX4*^−/−^ HT1080 cells expressing wild-type GPX4, treatment with the FGFR1 inhibitor PD166866 markedly increased ferroptosis and lipid peroxidation; however, in cells expressing the 2YF, 2YD or 2YE mutants, FGFR1 inhibition did not further aggravate these phenotypes (**Fig. 4k**). Similar results were observed in primary mouse cardiomyocytes subjected to H/R (**Fig. 4l**). These data establish that FGFR1-mediated phosphorylation of GPX4 at Tyr180/Tyr196 is a critical determinant of ferroptosis sensitivity.

### Activation of the FGFR1-GPX4 axis suppresses ferroptosis-mediated cardiac I/R injury

Given that ferroptosis has been identified as a major driver of myocardial I/R injury^8,29^, we investigated the functional role of the FGFR1-GPX4 axis in this pathological context. In heart tissue subjected to I/R, tyrosine phosphorylation of GPX4 progressively declined, paralleling reductions in activated FGFR1 (p-Tyr653/654) levels (**Extended Data Fig. 10a**), suggesting that this signaling axis is suppressed during I/R injury. To elucidate the role of GPX4 in cardiac I/R pathology, we generated cardiomyocyte-specific *Gpx4* knockout (*Gpx4*-CKO) mice (**Extended Data Fig. 10b-d**). Compared with *Gpx4*^f/f^ controls, *Gpx4*-CKO mice exhibited significantly elevated levels of MDA and canonical ferroptosis markers (*Ptgs2* mRNA and 4-HNE protein) after I/R (**Extended Data Fig. 10e-h**), indicating markedly enhanced ferroptosis. *Gpx4*-CKO mice also showed exacerbated cardiac injury, as evidenced by increased serum levels of cardiac injury markers (CK-MB, AST and LDH), upregulation of heart failure marker mRNAs (*Anp* and *Bnp*), impaired contractile function assessed by echocardiography, and significantly enlarged infarct size (**Extended Data Fig. 10i-n**). Collectively, these findings establish GPX4 as a critical endogenous suppressor of ferroptosis in the heart. Its loss exacerbates I/R injury, suggesting that its activation could confer protection.

We previously demonstrated that the natural FGFR1 ligand FGF-1 inhibits I/R-induced ferroptosis through FGFR1 activation (**Fig. 2k, l**). However, FGF-1 binds to and activates all four FGFR subtypes with limited selectivity^37,38^. Structural analysis revealed that FGF-1 interacts with FGFR2-4 via its N-terminal domain, whereas the FGF-1 region that interacts with FGFR1 lies outside this N-terminal region (**Extended Data Fig. 11a**). Based on this difference, we engineered an N-terminally truncated recombinant ligand, FGF-1^ΔNT^, that lacks the first 24 amino acids (**Extended Data Fig. 11b-d**). SPR assays showed that FGF-1^ΔNT^ exhibited markedly reduced affinity for FGFR2-4 while retaining normal binding to FGFR1 (**Extended Data Fig. 11e, f**). Functionally, FGF-1^ΔNT^ selectively activated FGFR1 signaling without inducing phosphorylation of FGFR2-4 (**Extended Data Fig. 11g-j**). In a mouse model of cardiac I/R, administration of FGF-1^ΔNT^ provided cardiac protection comparable to that of wild-type FGF1 (FGF-1^WT^), as evidenced by attenuated myocardial injury, preserved ventricular contractile function, reduced infarct size, and suppression of ferroptosis markers (*Ptgs2,* 4-HNE, and oxidized lipids) (**Extended Data Fig. 12a-h**). These results indicate that FGF-1^ΔNT^ is a functionally selective FGFR1 agonist with therapeutic potential for mitigating cardiac I/R injury through ferroptosis modulation.

To determine whether FGFR1-mediated tyrosine phosphorylation of GPX4 is functionally required for cardiac protection against I/R injury, we used adeno-associated virus (AAV) to express either wild-type GPX4 (WT-GPX4) or a phosphorylation-deficient mutant (2YF-GPX4) in the hearts of cardiomyocyte-specific *Gpx4* knockout (*Gpx4*-CKO) mice (**Fig. 5a, b**). Under I/R stress, mice expressing 2YF-GPX4 exhibited exacerbated myocardial injury, left ventricular dysfunction and enlarged infarct size compared with mice expressing WT-GPX4 (**Fig. 5c-h**). Importantly, the cardioprotective effects of FGF-1^ΔNT^, including the reduction of serum CK-MB, AST and LDH levels, preservation of cardiac function, and suppression of hypertrophy markers (*Anp* and *Bnp*), were abrogated in mice expressing 2YF-GPX4 (**Fig. 5c-h**). Consistent with this result, the anti-ferroptotic effect of FGF-1^ΔNT^, reflected by decreased levels of MDA, PTGS2, 4-HNE and oxidized lipids in the heart, was prominent in hearts reconstituted with WT-GPX4, but absent in those expressing 2YF-GPX4 (**Fig. 5i-l**). Therefore, our findings establish FGFR1-dependent tyrosine phosphorylation of GPX4 at Tyr180/Tyr196 as a critical regulatory mechanism that protects the heart against I/R injury by suppressing ferroptosis. Selective activation of the FGFR1-GPX4 axis, for instance using FGF-1^ΔNT^, represents a potential therapeutic strategy for ischemic heart disease.

**Fig. 5.**
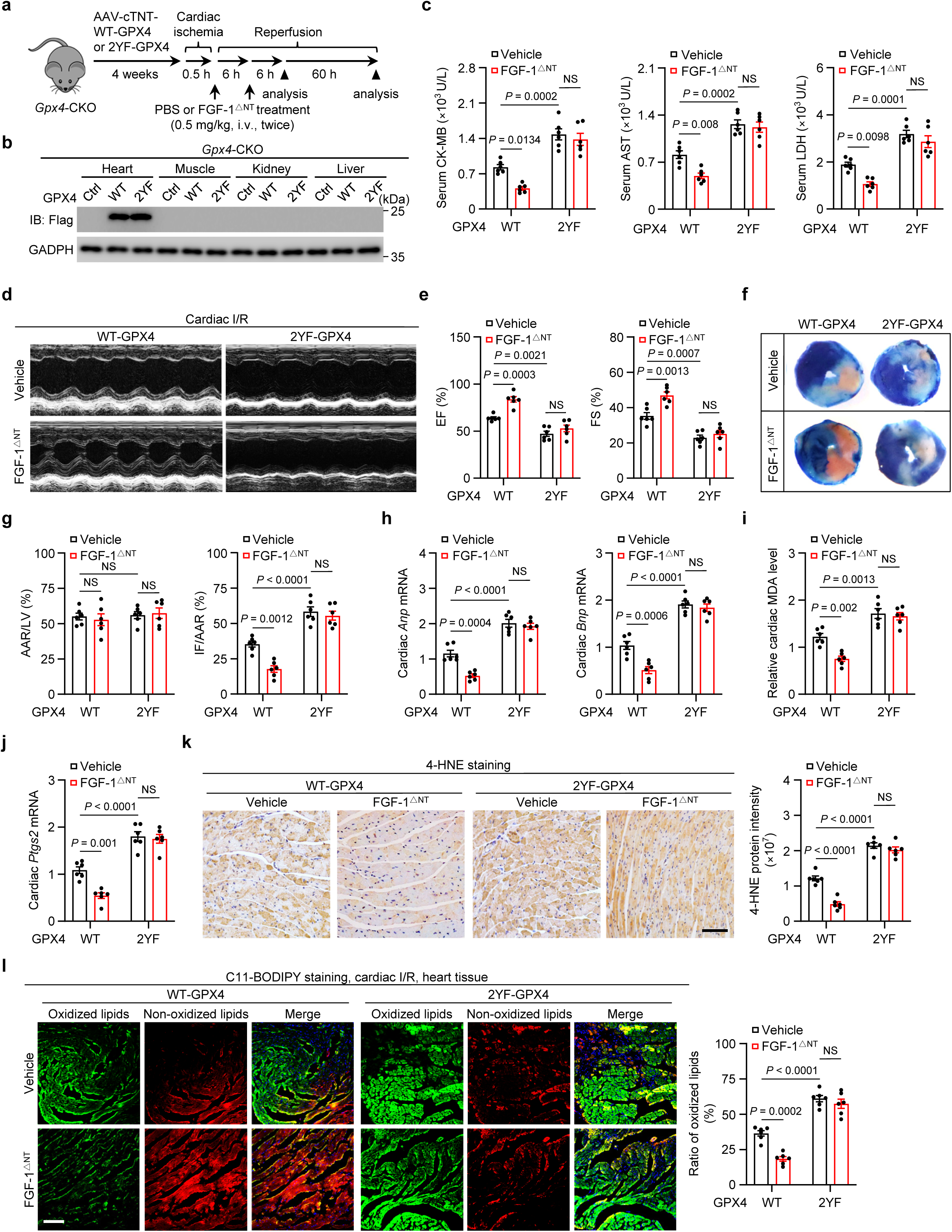
FGFR1 mediated GPX4 phosphorylation modulates cardiac I/R injury via ferroptosis. **a,** Schematic of treatment regimen with FGF-1^ΔNT^ in mice with cardiac I/R injury. The cardiomyocyte-specific *Gpx4* knockout (*Gpx4*-CKO) mice expressing either WT-GPX4 or 2YF-GPX4 for 4 weeks, followed by treatment with PBS (Vehicle) or FGF-1^ΔNT^ (0.5 mg/kg body weight, i.v. twice). **b,** Immunoblot analysis of GPX4 in tissue lysates (heart, muscle, kidney, and liver) from cardiac-specific *Gpx4* deficiency (*Gpx4*-CKO) mice expressing either WT-GPX4 or 2YF-GPX4 for 4 weeks. **c,** Serum CK-MB, AST, and LDH levels in the indicated mice subjected to cardiac I/R injury. **d,e,** Representative echocardiography images (**d**) and statistical analysis (**e**) of ejection fraction (EF) and fractional shortening (FS) in the indicated mice after cardiac I/R injury. **f,g,** Representative images (**f**) and quantitative data (**g**) for relative area at risk (AAR) and infarct size (IF) in heart sections from the indicated mice. **h,** Relative mRNA levels of cardiac *Anp* and *Bnp* in the indicated mice. **i,** Relative cardiac malondialdehyde (MDA) levels in the indicated mice after cardiac I/R injury. **j,** Relative *Ptgs2* mRNA in heart from the indicated mice. **k,** Representative images and quantification of 4-HNE–stained cardiac sections from the indicated mice. Scale bar, 100 μm. **l,** Representative images and quantification of C11-BODIPY staining in heart sections from the indicated groups. Scale bar, 100 μm. (**c, e, g-l**) *n* = 6 mice/group. Data are mean ± s.e.m.; ordinary two-way ANOVA, followed by Tukey in (**c, e, g-j,** right graphs of **k** and **l**); \**p* < 0.05, \*\**p* < 0.01, \*\*\**p* < 0.001, \*\*\*\**p* < 0.0001, NS, not significant; Gel source data blots can be found in Supplementary Fig. 1.

### Cardiomyocyte-specific *Gpx4*-Y180/196F knock-in mice are hypersensitive to cardiac I/R injury via ferroptosis

To further validate the functional importance of FGFR1-mediated GPX4 phosphorylation under physiological expression levels, we generated a cardiomyocyte-specific conditional knock-in mouse model in which the endogenous wild-type *Gpx4* allele was replaced with 2YF-*Gpx4* (*Gpx4*^2YF^-KI-CKO), the phosphorylation-deficient version (**Extended Data Fig. 13a-c**). These *Gpx4*^2YF^-KI-CKO mice were viable and showed no overt developmental abnormalities. Under I/R stress, *Gpx4*^2YF^-KI-CKO mice exhibited significantly elevated levels of 4-HNE, MDA and the ferroptosis marker Ptgs2 and oxidized lipids in the heart compared with control *Gpx4*^2YF^-KI^f/f^ littermates (**Fig. 6a-e**), indicating exacerbated cardiac ferroptosis. Consistent with the pathophysiological link between ferroptosis and I/R injury, *Gpx4*^2YF^-KI-CKO mice developed more severe myocardial injury, impaired left ventricular contractile function and enlarged infarct size (**Fig. 6f-k**). Notably, the cardioprotective effects of the selective FGFR1 agonist FGF-1^ΔNT^ were abrogated in *Gpx4*^2YF^-KI-CKO mice (**Fig. 6f-k**). Furthermore, we investigated the long-term impact of this axis on I/R injury. We found that the 2YF-*Gpx4* mutation promoted I/R-induced cardiac remodeling and fibrosis, accompanied by the upregulation of the profibrotic gene Col1α1 (**Extended Data Fig. 13d-f**). FGF-1^△NT^ treatment significantly attenuated fibrosis in *Gpx4*^2YF^-KI^f/f^ mice but failed to exert this therapeutic effect in *Gpx4*^2YF^-KI-CKO mice (**Extended Data Fig. 13d-f**). Collectively, these findings corroborate that the cardioprotective effects of FGF-1^ΔNT^ specifically require FGFR1-mediated phosphorylation of GPX4 at Tyr180 and Tyr196.

**Fig. 6.**
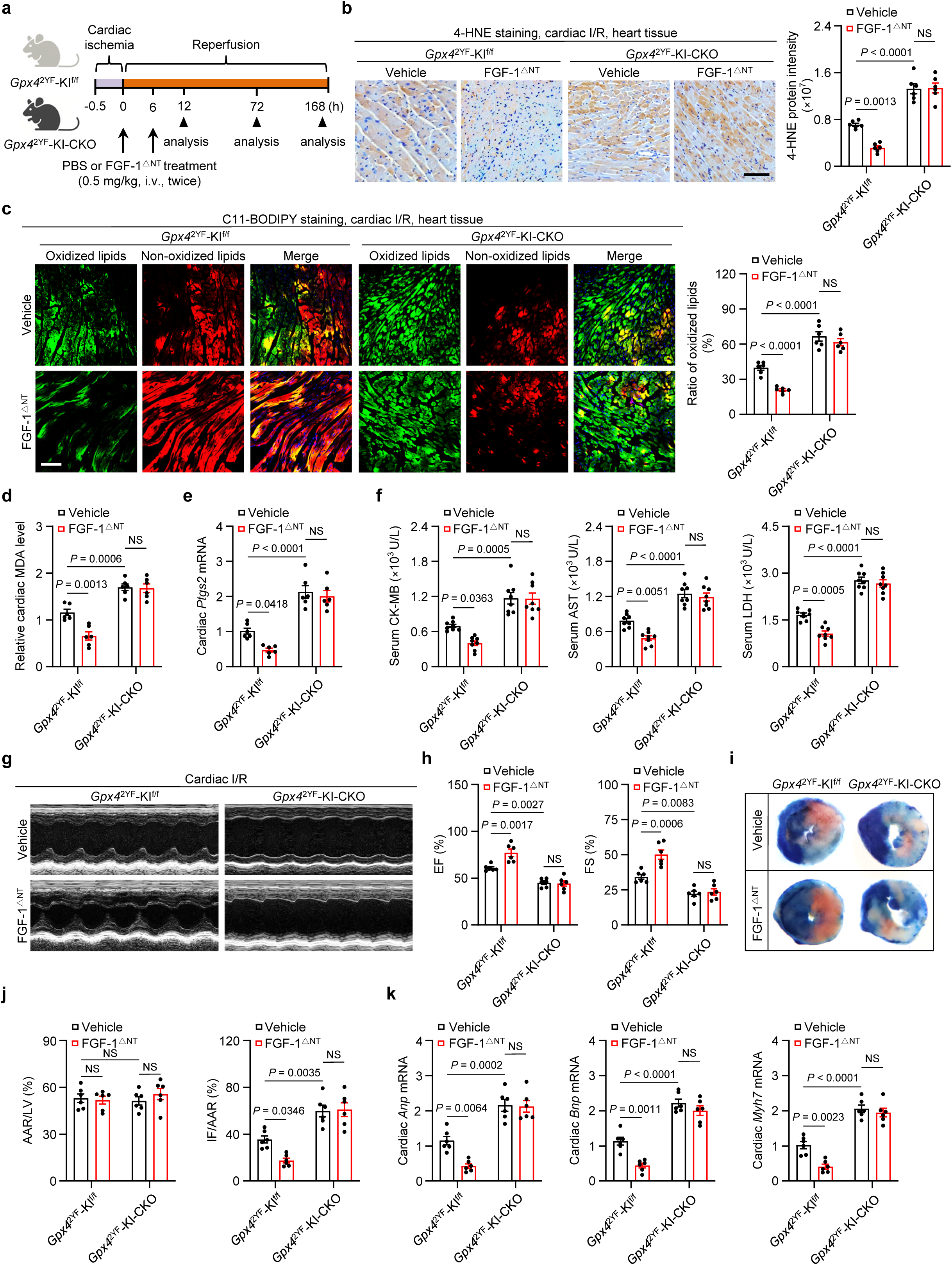
GPX4-Y180/196F knock-in mice exhibit hypersensitivity to cardiac I/R injury. **a,** Treatment regimen for FGF-1^ΔNT^ in *Gpx4*-Y180/196F-knock-in-flox/flox (*Gpx4*^2YF^-KI^f/f^) and cardiomyocyte-specific *Gpx4*-Y180/196F-knock-in (*Gpx4*^2YF^-KI-CKO) mice subjected to cardiac I/R injury. Mice received either PBS (Vehicle) or FGF-1^ΔNT^ (0.5 mg/kg, i.v., twice). **b,** Representative images and quantification of 4-HNE–stained cardiac sections from the indicated mice. Scale bar, 100 μm. **c,** Representative images and quantification of C11-BODIPY staining in heart sections from the indicated groups. Scale bar, 100 μm. **d,** Relative cardiac MDA levels in the indicated mice. **e,** Cardiac *Ptgs2* mRNA in the indicated mice. **f,** Serum CK-MB, AST and LDH levels in indicated mice after cardiac I/R injury. *n* = 8 mice/group. **g,h,** Representative echocardiography images (**g**) and statistical analysis (**h**) of EF and FS in indicated mice. **i,j,** Representative images (**i**) and quantitative data (**j**) for relative AAR and IF in heart section obtained from indicated mice. **k,** Relative mRNA levels of cardiac *Anp*, *Bnp* and *Myh7* in indicated mice. (**b-e, h, j, k**) *n* = 6 mice/group. Data are mean ± s.e.m.; ordinary two-way ANOVA, followed by Tukey in (**d-f, h, j, k,** right graph of **b** and **c**); \**p* < 0.05, \*\**p* < 0.01, \*\*\**p* < 0.001, \*\*\*\**p* < 0.0001, NS, not significant.

To determine whether the exacerbated injury phenotype could be rescued by pharmacological inhibition of ferroptosis, we pretreated *Gpx4*^2YF^-KI^f/f^ and *Gpx4*^2YF^-KI-CKO mice with the ferroptosis inhibitor Fer-1 before I/R (**Extended Data Fig. 14a**). Fer-1 administration effectively suppressed ferroptotic signaling, as evidenced by the reduced accumulation of 4-HNE and PTGS2 (**Extended Data Fig. 14b-d**), and significantly ameliorated I/R-induced cardiac dysfunction in both genotypes (**Extended Data Fig. 14e-j**). In addition, Fer-1 treatment attenuated interstitial fibrosis in both groups (**Extended Data Fig. 14k, l**). Importantly, Fer-1 rescued the hypersensitivity to I/R injury conferred by the 2YF-*Gpx4* mutation (**Extended Data Fig. 14e-l**).

Together, these results establish phosphorylation of GPX4 at Tyr180 and Tyr196 as a critical molecular switch that protects the heart against I/R injury. This phosphorylation event serves as the functional bridge linking FGFR1 signaling to GPX4; its loss renders the heart highly susceptible to ferroptosis-mediated damage, a vulnerability that can be effectively overcome by therapeutic targeting of ferroptosis.

## Discussion

In this study, we identify a previously unrecognized RTK-dependent tyrosine phosphorylation axis in which the FGFR1 directly phosphorylates the core ferroptosis regulator GPX4 at Tyr180 and Tyr196, thereby enhancing GPX4 enzymatic activity and suppressing ferroptosis. This finding challenges the prevailing view that levels of GPX4 activity are regulated via transcriptional control or cofactor availability^4,14,18,39,40^, establishing tyrosine phosphorylation of GPX4 as a rapid-response regulatory mechanism. Moreover, the direct interaction between FGFR1 and GPX4 functionally links RTK signaling to the ferroptosis machinery – two areas broadly implicated in physiology and disease but not previously connected. We therefore propose the concept of a “phosphorylation switch”, wherein GPX4 acts as a redox sensor that integrates extracellular stimuli with intracellular lipid peroxidation stress by receiving phosphorylation signals from a growth factor receptor.

Unlike canonical FGFR signaling, which engages multiple downstream effectors including the adaptor protein FRS2 and the lipase PLCγ^32,33^, FGFR1 directly binds and phosphorylates GPX4, representing a non-canonical mode of signal transduction. Domain mapping revealed that the kinase domain of FGFR1 engages GPX4, with Tyr730 of FGFR1 serving as the critical binding residue. This interface is highly evolutionarily conserved. Through site-directed mutagenesis, phospho-specific antibodies, and enzymatic activity assays, we unambiguously demonstrate that FGFR1 phosphorylates Tyr180 and Tyr196 within the GPX4 protein. This phosphorylation enhances GPX4 catalytic activity by increasing its binding affinity for lipid peroxides, rather than by altering GSH levels or protein stability. Notably, phosphomimetic GPX4 mutants (2YD/2YE) are not constitutively active, indicating that the structural effect of tyrosine phosphorylation is highly specific and cannot be recapitulated by simply introducing a charge. Furthermore, both the binding and phosphorylation of GPX4 by FGFR1 strictly depend on FGFR1 kinase activity and require Tyr730 as a docking site. We thus propose an alternative kinase-activity-dependent substrate recognition model in which FGFR1 autophosphorylation at Tyr730 creates a dedicated docking interface for GPX4, enabling subsequent phosphorylation of GPX4 at Tyr180/Tyr196 to enhance its catalytic activity and suppress ferroptosis.

Previous studies have largely focused on transcriptional regulation (e.g., NRF2)^17,18,41^, protein stability (e.g., deubiquitinases or autophagy)^19–21,42–45^, and GSH availability to control the levels of GPX4 activity^14,22,40^. Our study introduces tyrosine phosphorylation as a rapid and distinct layer of regulation. Rather than conflicting with the classical paradigm of GPX4 inactivation leading to ferroptosis^14^, GPX4 phosphorylation provides an upstream activation signal that enables rapid cellular responses to extracellular cues, such as growth factors, enhancing resistance to ferroptosis. Compared with other ferroptosis regulators^46–51^, FGFR1, as an RTK, offers the unique advantage of being selectively activatable by ligands, making it a tractable drug target. Moreover, the antiferroptotic effect of FGFR1 is cell-type-dependent – only seen in cells that express high levels of FGFR1 – suggesting that FGFR1 could serve as a critical determinant of ferroptosis susceptibility and enable personalized therapeutic strategies in ischemic disease.

In the context of myocardial I/R injury, FGFR1 activity (p-Tyr653/654) and GPX4 phosphorylation levels decline in parallel and are inversely correlated with ferroptosis markers, indicating that this axis is suppressed under pathological conditions. Notably, lipid peroxidation itself can further suppress FGFR1 activity. This establishes a positive-feedback loop in which ferroptosis leads to FGFR1 inactivation, which in turn promotes GPX4 dephosphorylation and amplifies ferroptosis. This loop provides a mechanistic explanation for the rapid propagation of cell death in I/R injury. Using structure-guided design^52,53^, we generated a selective FGFR1 agonist, FGF-1^ΔNT^, which avoids the off-target effects associated with activating FGFR2-4. This agonist attenuates I/R injury in wild-type mice but fails to protect mice expressing a phosphorylation-deficient version of GPX4 (*Gpx4*^2YF^-KI-CKO), validating that GPX4 phosphorylation is an essential downstream effector. Furthermore, the ferroptosis inhibitor Fer-1 fully rescues the hypersensitive phenotype of *Gpx4*^2YF^-KI-CKO mice, demonstrating that the exacerbated injury in these mice is indeed mediated by ferroptosis and that direct targeting of ferroptosis remains effective even when GPX4 phosphorylation is compromised.

Taken together, our findings connect two seemingly independent fields – RTK signaling and ferroptosis – through the direct interaction and phosphorylation of GPX4 by FGFR1. This discovery expands the network of ferroptosis regulators and reveals new therapeutic strategies for treating ischemic heart disease and other ferroptosis-associated pathologies, namely through selective activation of the FGFR1-GPX4 axis or direct inhibition of ferroptosis. These two approaches may be complementary, enabling precision interventions based on patient molecular stratification. From a broader biological perspective, GPX4, acting as a redox sensor that receives phosphorylation signals from a growth factor receptor, exemplifies how cells integrate metabolic adaptation (antioxidant defense) with proliferation/survival signals in response to a changing environment.

## Methods

### Mouse models

All animal procedures were approved by the Institutional Animal Care and Use Committee (IACUC) at Wenzhou Medical University and conducted in strict compliance with ethical guidelines, under approval number SYXK2020-0014. Mice were housed in a temperature-controlled environment under a 12-h light/dark cycle, with *ad libitum* access to water and standard rodent chow. Male C57BL/6J mice were obtained from Beijing Vital River Laboratory Animal Technology Co., Ltd. *Myh6*-Cre mice (Cat. C001041) was purchased from Cyagen Biosciences Inc. *Gpx4*-Y180/196F knock-in (*Gpx4*^2YF^*-*KI^f/f^) mice were generated from Cyagen Biosciences Inc. *Fgfr1*^flox/flox^ (*Fgfr1*^f/f^) mice were kindly provided by Prof. Li Lin (Wenzhou Medical University), and *Gpx4*^flox/flox^ (*Gpx4*^f/f^) mice were provided by Prof. Fudi Wang (Zhejiang University). Cardiomyocyte-specific knockout (*Fgfr1*-CKO or *Gpx4*-CKO) or knock-in (*Gpx4*^2YF^-KI-CKO) mice were generated by crossing *Fgfr1*^f/f^, *Gpx4*^f/f^, or *Gpx4*^2YF^-KI^f/f^ mice with *Myh6*-Cre transgenic mice, respectively. Offspring genotype was performed by PCR amplification of genomic DNA extracted from tail biopsies, using primers listed in **Supplementary Table S1**.

To generate *Gpx4*^2YF^-KI^f/f^ mice, a targeting strategy was designed to introduce conditional knock-in mutations into the *Gpx4* gene. This strategy employed two loxP sites flanking a region containing two tyrosine-to-phenylalanine substitutions (p.Y180F and p.Y196F; TAT to TTC for both sites). Briefly, the donor vector was constructed to contain the following cassette: loxP-endogenous splice acceptor of intron 4-Exon 5 to Exon 7 of mouse *Atpif1*-6xSV40 pA-loxP-KI region (harboring the p.Y180F and p.Y196F mutations). The gRNA targeting the mouse *Gpx4* gene, the linearized donor vector, and Cas9 mRNA were co-injected into fertilized mouse eggs to generate targeted conditional knock-in offspring. The injected embryos were transferred into pseudopregnant foster mothers. F0 founder animals were identified by PCR genotyping followed by Sanger sequence analysis to confirm the correct integration of the knock-in cassette and the presence of the desired point mutations. The positive F0 founders were then bred to wild-type C57BL/6J mice to test germline transmission and establish the F1 generation. The genotypes of the resulting progeny were determined by PCR analysis using the following specific primers: primer F, 5’-GCAAGTCTGTGTCATGCATGCAT-3’; primer R, 5’-GGCAAGTGGATATGAGAGTTCTAAGC-3’. The germline-transmitted F1 mice were further backcrossed to the C57BL/6J background for subsequent experiments.

For AAV-mediated cardiomyocyte-specific *Gpx4* overexpression, 8-week-old *Gpx4*-CKO mice were intravenously injected with 100 µL of virus suspension containing 5 × 10^10^ vector genomes of either AAV2/9-cTNT-3×Flag-WT-GPX4 or AAV2/9-cTNT-3×Flag-GPX4-Y180/196F (AAV2/9-cTNT-3×Flag-2YF-GPX4) via tail vein. Four weeks post-infection, mice were subjected to cardiac ischemia-reperfusion surgery.

### Antibodies and drugs

The polyclonal antibodies specifically recognizing phosphorylated GPX4 were generated by immunizing rabbits with the following synthetic peptides: VKR (pTyr) GPMEE for phospho-Tyr180-GPX4 and EKDLPH (pTyr) FC for phospho-Tyr196-GPX4. Antibodies to FGF Receptor 1 (1:1000 for IB, 1:50 for IP, Cat. 9740), phospho-FGF Receptor 1 (Tyr653/654) (1:1000, Cat. 52928), phospho-Tyrosine (1:2000, Cat. 8954), c-Casp3 (1:1000, Cat. 9661), PARP (1:1000, Cat. 9542), c-Casp1 (1:1000, Cat. 89332), Casp1 (1:1000, Cat. 24232), PLCγ (1:1000, Cat. 5690), Col1a1 (1:200 for IHC, Cat. 72026), Normal IgG (1:50 for IP, Cat. 2729), phospho-FGF Receptor (Tyr653/654) (1:1000, Cat. 3471), FGFR2 (1:1000, Cat. 23328) and FGFR3 (1:1000, Cat. 4574), phospho-ERK1/2 (Thr202/Tyr204) (1:2000, Cat. 4370) and ERK1/2 (1:2000, Cat. 9102) were purchased from Cell Signaling Technology. Antibodies to Flag (1:2000, Cat. 20543-1-AP), HA (1:5000, Cat. 51064-2-AP), RIP3 (1:1000, Cat. 17563-1-AP), MLKL (1:1000, Cat. 66675-1-AP), FRS2 (1:1000, Cat. 11503-1-AP), GAPDH (1:1000, Cat. 60004-1-Ig), His-Tag (1:5000, Cat. 66005-1-Ig), GST-tag (1:10000, Cat. 66001-2-Ig) and FGFR4 (1:1000, Cat. 11098-1-AP) were obtained from Proteintech. Antibodies to phospho-RIP3 (1:1000, Cat. ab222320), phospho-MLKL (1:1000, Cat. ab196436), NT-GSDMD (1:1000, Cat. ab215203), GSDMD (1:1000, Cat. ab219800) and 4-HNE (1:50 for IHC, Cat. ab48506) were purchased from Abcam. Antibodies to HA (F7 for IP, 1:100, Cat. sc-7392; Y11 for IB, 1:500, Cat. sc-805) were obtained from Santa Cruz Biotechnology. Anti-GPX4 (1:5000 for IB, 1:100 for IP, Cat. A11243), β-actin (1:20000, Cat. A5441), Flag (1:100 for PLA, Cat. MA1-91878) and Casp3 (1:1000, Cat. DF6879) was purchased separately from ABclonal, Thermo Fisher Scientific, Sigma-Aldrich and Affbiotech. Goat anti-rabbit IgG-HRP (1:10000, Cat. 111-035-003) and goat anti-mouse IgG-HRP (1:10000, Cat. BS12478) were from Jackson ImmunoResearch Labs and Bioworld Technology, respectively.

Anti-Flag beads (M2) (Cat. M8823), 3 × Flag peptide (Cat. F3290), LPS (Cat. L4516), GSH (Cat. G4251), glutathione reductase (GR) (Cat. G3664), NADPH (Cat. N7505), and Protease XIV (Cat. P5147) and tert-butyl hydroperoxide (TBHP) (Cat. 458139) were purchased from Sigma-Aldrich. Collagenase II (Cat. LS004176) and Collagenase IV (Cat. LS004188) were purchased from Worthington Biochemical. FGFR1 inhibitors PD166866 (Cat. S8493) and PD173074 (Cat. S1264), Fer-1 (Cat. S7243), DFO (Cat. S5742),

Necrostatin-1 (Cat. S8037), Lip-1 (Cat. S7699), RSL3 (Cat. S8155), Erastin (Cat. S7242) and Blasticidin (Cat. S7419) were purchased from Selleck. Staurosporine (STS, Cat. HY-15141), cycloheximide (CHX, Cat. HY-12320), SM-164 (Cat. HY-15989A), Z-VAD-FMK (Cat. HY-16658) and Nigericin (Cat. HY-127019) were purchased from MedChemExpress. TNFα (Cat. CF09) was obtained from Novoprotein. C11-BODIPY (581/591) (Cat. D3861) was purchased from Thermo Fisher Scientific. PCOOH (Cat. 31976) was purchased from Cayman.

### Generation of CRISPR/Cas9-mediated knockout cell lines

CRISPR-Cas9-mediated knockout of *FGFR1* and *GPX4* in HT1080 cells, as well as *Fgfr1*, *Plcγ* or *Frs2* knockout in MEFs, were performed as described^54^. The sequence for each sgRNA was: 5’-TTTTCAACCAGCGCAGTGTG-3’ (sg1) and 5’-GATCCGGTCAAATAATGCCT-3’ (sg2) for human *FGFR1*; 5’-CACGCCCGATACGCTGAGTG-3’ (sg2) for human *GPX4*; 5’-CTCTCATGAGCCGTTACCTG-3’ (sg1) and 5’-TCTCCGAGATCAGATCCGAC-3’ (sg2) for mouse *Fgfr1*; 5’-GCTACGCTACCCCATCAACG-3’ (sg1) and 5’-GCGGCGCGGACAAAATCGAG-3’ (sg2) for mouse *Plcγ*; 5’-CGAGATGTACCGTCGTCTCG-3’ (sg1) and 5’-AGTCATAGCCGTATCGTCGT-3’ (sg2) for mouse *Frs2*. To construct the knockout cell lines, cells were co-transfected with 1.5 μg of gRNA-expressing plasmid and 1.5 μg of human codon-optimized Cas9 (hCas9) plasmid, followed by blasticidin (Selleck, Cat. S7419) selection for 3 days. Single-cell clones were expanded, isolated, and validated by DNA sequencing and immunoblotting. Primer sequences are also shown in **Supplementary Table S2**. Because *GPX4* loss alone is sufficient to trigger ferroptosis, *GPX4* ^−/−^HT1080 cell were generated and propagated in the presence of the ferroptosis inhibitor liproxstatin-1 (Lip-1), which was removed for testing various assays throughout the study.

### Generation of the lentiviral system

Lentiviral particles were generated by transfecting HEK293T cells with a lentiviral vector and packaging plasmids using a liposomal transfection reagent (YEASEN, Cat. 40802ES03). At 48 h post-transfection, the culture supernatants were harvested, filtered through a 0.45 μm filter, and stored at −80 °C until further use. For infection, the cells were cultured in virus-containing medium supplemented with 10 μg/mL polybrene. To enhance viral transduction, the plates were centrifuged at 1,000 *g* for 30 min. The medium was replaced 12 h after infection. All experiments were performed at least 48 h post-infection.

### Cell viability and death assays

For cell viability assays, cells were seeded in 96-well plates at a density of 3,000 cells/well. After 24 h of incubation, cells were treated as indicated and subsequently incubated with Cell Counting Kit-8 (CCK-8; DOJINDO, Cat. CK04) for 2 h at 37 °C with 5% CO_2_. Prior to measurement, plates were gently agitated on an orbital shaker for 1 min to ensure a homogeneous color distribution. Absorbance was measured at 450 nm using a microplate reader.

For cell death analysis, cells were seeded in 6-well plates and allowed to reach ∼50% confluency. After 24 h, cells were subjected to the indicated treatments for various durations. Both adherent and floating cells were collected and stained with 5 μg/mL propidium iodide (PI; Vazyme, Cat. A211-02) for quantification by flow cytometry. A minimum of 10,000 cells were acquired per sample, and data analysis was performed using FlowJo software. All experiments were conducted in at least triplicate.

### Lipid peroxidation assay

Cells were seeded in 6-well plates at a density of 1.5 × 10^5^ cells/well and incubated overnight at 37 °C with 5% CO_2_. Following treatment, cells were incubated with 1 μM C11-BODIPY (581/591) (Thermo Fisher Scientific, Cat. D3861) for 30 min at 37 °C. Subsequently, cells were harvested by trypsinization, resuspended in PBS, and filtered through a 40 μm cell strainer (FALCON, Cat. 352340). Lipid ROS levels were quantified by flow cytometry using 488 nm excitation and FL1 detection. A minimum of 10,000 single-cell events were acquired per sample, and data were analyzed using FlowJo software. For tissue analysis, frozen heart tissue sections (5 μm) were equilibrated to room temperature, rinsed with PBS, then stained with 5 μM C11-BODIPY (581/591) for 10 min at 37 °C. Confocal microscopy was used to detect the oxidized lipid (green) and non-oxidized lipid (red) signal.

### Cell culture and treatment

Mouse embryonic fibroblasts (MEFs) were established by infecting primary MEFs harvested at embryonic day 13.5, with a lentivirus encoding SV40 large T antigen. U251 and U87 cells were kindly provided by Prof. Yin Chen (Xiamen University). HS-578T, NCI-H1299, HCT116, HeLa, NCI-H1975, NCI-H1573, HEK293T and NCI-H209 cells were gifts from Prof. Sheng-Cai Lin (Xiamen University). MDA-MB-231, Huh7, HepG2 and Hep3B cells were provided by Prof. Qiao Wu (Xiamen University). The human fibrosarcoma cell line HT1080 (Cat. CC2011) was purchased from CellCOOK. L6 rat skeletal myoblasts ectopically expressing human FGFR1, FGFR2, FGFR3, or FGFR4 were generously provided by Prof. Gaozhi Chen (Wenzhou Medical University). All cell lines were confirmed to be mycoplasma-free, and HT1080 cells were authenticated by short tandem repeat (STR) profiling.

All cells were maintained at 37 °C in a humidified atmosphere containing 5% CO_2_. U251, U87, HEK293T, HS-578T, NCI-H1299, HCT116, HeLa, NCI-H1975, MEF, L6 rat skeletal myoblasts, MDA-MB-231, HepG2, Huh7, and Hep3B cells were cultured in Dulbecco’s Modified Eagle’s Medium (DMEM; Gibco, Cat. 11995065) supplemented with 10% fetal bovine serum (FBS), 100 IU/mL penicillin, and 100 μg/mL streptomycin (P/S). HT1080 cells were cultured in Minimum Essential Medium (MEM; Gibco, Cat. 41090036) supplemented with 10% FBS and P/S. NCI-H1573 and NCI-H209 cells were maintained in RPMI-1640 medium (Gibco, Cat. 11875093) supplemented with 10% FBS and P/S.

For apoptosis induction, MEFs were treated with 0.5 μM Staurosporine (STS; MedChemExpress, Cat. HY-15141) for the indicated times. Alternatively, cells were pretreated with 1 μg/mL cycloheximide (CHX; MedChemExpress, Cat. HY-12320) for 30 min, followed by co-treatment with 10 ng/mL TNFα (Novoprotein, Cat. CF09) and CHX. Necroptosis was triggered by treating cells with a TSZ cocktail consisting of 10 ng/mL TNFα, 0.1 μM SM-164 (MedChemExpress, Cat. HY-15989A), and 20 μM Z-VAD-FMK (MedChemExpress, Cat. HY-16658) for the indicated durations. To induce pyroptosis, cells were primed with 1 µg/mL lipopolysaccharide (LPS; Sigma-Aldrich, Cat. L4516) for 5 h, followed by stimulation with 20 µM Nigericin (MedChemExpress, Cat. HY-127019). To induce ferroptosis, cells were treated with RSL3 (Selleck, Cat. S8155) or Erastin (Selleck, Cat. S7242) at the indicated concentrations for indicated durations. To investigate the role of FGFR1 in ferroptosis, cells were treated with 20 μM PD166866 (Selleck, Cat. S8493) or 50 nM PD173074 (Selleck, Cat. S1264) as indicated.

For cystine starvation assays, MEFs were washed twice with PBS and cultured in a specific starvation medium for 9 h. This medium consisted of glutamine-, methionine-, and cystine-free DMEM (Gibco, Cat. 21013024) supplemented with 10% dialyzed FBS (Bioexplorer, Cat. BD1643-015), 100 μg/mL P/S, 2 mM L-Glutamine (Gibco, Cat. 25030081), and 0.1 mM L-Methionine (Sigma-Aldrich, Cat. M9625). For the control group, the medium was further supplemented with 0.1 mM L-Cystine (Sigma-Aldrich, Cat. C8755).

Pharmacological rescue experiments were performed by co-treating cells with the following inhibitors: ferroptosis inhibitors ferrostatin-1 (Fer-1, 5 μM; Selleck, Cat. S7243), liproxstatin-1 (Lip-1, 5 μM; Selleck, Cat. S7699) and deferoxamine mesylate (DFO, 5 μM; Selleck, Cat. S5742), the necroptosis inhibitor necrostatin-1 (Nec1, 2 μM; Selleck, Cat. S8037), or the pan-caspase inhibitor Z-VAD-FMK (ZVF, 10 μM; MedChemExpress, Cat. HY-16658).

### Cardiac ischemia-reperfusion (I/R) model

To induce I/R injury, male mice (8-10-week-old) were anesthetized with isoflurane, intubated, and mechanically ventilated. Body temperature was maintained at 37 °C using a heating pad. A left thoracotomy was performed at the third intercostal space, and the left anterior descending (LAD) coronary artery was ligated with a 7-0 silk suture. Ischemia was confirmed by immediate blanching of the anterior left ventricular wall. After 30 min of ischemia, the slipknot was released to allow reperfusion, which was verified by the restoration of myocardial color. Sham-operated mice underwent an identical procedure without LAD ligation. For drug treatments, mice received Fer-1 (1.0 mg/kg, i.p.) 24 h and 1 h before surgery. Wild-type FGF-1 (FGF-1^WT^, 0.5 mg/kg) or the N-terminally truncated FGF-1 variant (FGF-1^ΔNT^, 0.5 mg/kg) was administered via tail vein injection immediately upon reperfusion (0 h) and at 6 h post-reperfusion. Mice were sacrificed at designated time points. Hearts were harvested, fixed overnight in 4% paraformaldehyde, paraffin-embedded, and sectioned at 5 µm thickness. Fibrosis was assessed using Masson’s Trichrome staining.

### Measurement of infarct size

Infarct size was quantified using the 2,3,5-triphenyltetrazolium chloride (TTC)-Evans blue double-staining technique. Briefly, the LAD was re-ligated under anesthesia, and Evans blue dye (1% w/v, Sigma-Aldrich, Cat. E2129) was immediately injected into the jugular vein to delineate the non-ischemic area. Hearts were excised, rinsed in PBS, and frozen at −20 °C for 15 min before being cut into 1-mm thick slices. Sections were incubated in 1% TTC solution (Sigma-Aldrich, Cat. T8877) at 37 °C for 30 min. Viable myocardium within the area at risk (AAR) stained red, while infarcted tissue remained pale white. Images were analyzed using ImageJ (NIH), and infarct size was expressed as a percentage of the AAR.

### Echocardiographic assessment

Cardiac function was evaluated using a high-resolution ultrasound system (Vevo 3100, VisualSonics Inc., Toronto, Canada) equipped with an MX400 linear array transducer (30 MHz). Mice were anesthetized with isoflurane 3 days post-surgery, maintaining a heart rate of ∼450 bpm. Two-dimensional parasternal long-axis images were acquired, followed by M-mode tracings perpendicular to the interventricular septum at the papillary muscle level. Left ventricular ejection fraction (EF%) and fractional shortening (FS%) were analyzed using Vevo LAB software based on standard M-mode measurements.

### Biochemical assays

Serum levels of creatine kinase MB isoenzyme (CK-MB; Cat. OSR61155), lactate dehydrogenase (LDH; Cat. OSR6128), and aspartate transaminase (AST; Cat. OSR6109) were quantified using a Beckman Coulter AU480 Chemistry Analyzer (Beckman Coulter Inc., Brea, CA, USA) according to the manufacturer’s instructions.

### Measurement of MDA and GSH content

Malondialdehyde (MDA) levels in cardiac tissue were measured using the MDA Assay Kit (Beyotime, Cat. S0131) in accordance with the manufacturer’s instructions. Glutathione (GSH) levels were determined using the GSH-Glo^TM^ Glutathione Assay kit (Promega, Cat. V6911), with luminescence measured on a Varioskan LUX microplate reader. Relative GSH levels were calculated by normalizing the luminescence intensity to the control group.

### Ferrous iron measurement

Ferrous iron (Fe^2+^) levels in cell lysates were quantified using an Iron Assay Kit (Dojindo, Cat. I291) according to a modified manufacturer’s protocol. A 2-fold serial dilution of the iron standard was prepared in assay buffer to generate a standard curve. To prevent non-specific reduction of ferric iron (Fe^3+^), the manufacturer-provided reducer solution was replaced with assay buffer (5% of the total sample volume). Following a 15 min pre-incubation at 37 °C, standards, cell lysate samples, and blank controls were transferred to a 96-well plate, and the probe solution was added to each well. The plate was incubated for 1 h at 37 °C, after which absorbance was measured at 593 nm using a microplate reader. Final Fe^2+^ concentrations were calculated from the standard curve and normalized to the initial sample dilution factor.

### Isolation of primary neonatal mouse cardiomyocytes

Primary neonatal mouse cardiomyocytes were isolated from *Fgfr1*-CKO neonatal mice using the Neonatal Heart Dissociation Kit (Miltenyi Biotec, Cat. 130-098-373). Excised ventricular tissues were minced and dissociated via sequential enzymatic digestion followed by mechanical trituration. The resulting single-cell suspension was pre-plated in 10-cm dishes for 1.5 h to deplete adherent cardiac fibroblasts. Non-adherent cardiomyocytes were then harvested and cultured in DMEM supplemented with 15% FBS. Adenoviral infection was performed 18–24 h after seeding, and ferroptosis assays were conducted 48 h post-infection.

### Primary mouse cardiomyocytes isolation and culture

Primary mouse cardiomyocytes were isolated using a modified injection-based *in situ* digestion method, as described^55^. Briefly, mice were anesthetized, and the heart was exposed. The ventricles were sequentially perfused via the apex with EDTA buffer to clear blood, followed by Perfusion Buffer, and finally Collagenase Buffer (containing Collagenase II, IV, and Protease XIV) for enzymatic digestion. The digested ventricles were manually dissociated into a single-cell suspension. Following filtration, viable cardiomyocytes were enriched by gravity sedimentation and subjected to a stepwise calcium reintroduction protocol (0.34 mM, 0.68 mM, and 1.02 mM Ca^2+^) to restore physiological calcium levels. Cells were then seeded onto Laminin-coated plates (5 µg/mL; Gibco, Cat. 23017015) in M199 (Sigma-Aldrich, Cat. M4530) culture medium. Unattached cells were removed after 1 h, and the medium was changed every 48 h.

### Hypoxia/reoxygenation (H/R) model

To mimic the I/R model *in vitro*, primary mouse cardiomyocytes were subjected to H/R. Briefly, cells were washed twice with PBS, and the M199 culture medium was replaced with glucose-free, serum-free DMEM (Gibco, Cat. 11966025) that had been pre-equilibrated in a hypoxic atmosphere. Cells were then incubated in a modular chamber (1% O_2_, 5% CO_2_, 94% N_2_) for 6 h at 37 °C to induce hypoxia. Following the hypoxic challenge, reoxygenation was initiated by replacing the starvation medium with fresh, complete M199 medium containing serum and antibiotics. Cells were then returned to normoxic conditions (21% O_2_, 5% CO_2_) for the indicated recovery times.

### Immunoprecipitation and immunoblotting

For immunoprecipitation (IP) of ectopically expressed proteins, cells seeded in 6-well dishes were transfected with the indicated expression plasmids using polyethylenimine (PEI; Polyscience, Cat. 23966). At 24 h post-transfection, cells were harvested in lysis buffer [20 mM Tris-HCl (pH 7.4), 150 mM NaCl, 1 mM EDTA, 1 mM EGTA, 1% Triton X-100, 2.5 mM sodium pyrophosphate, 1 mM β-glycerophosphate, 1 mM Na_3_VO_4_, 2 μg/mL leupeptin, and 1 mM PMSF], followed by sonication and centrifugation at 12,000 rpm for 10 min at 4 °C. An aliquot of the supernatant was reserved as the whole cell lysate. The remaining supernatant was incubated with anti-HA or Flag antibodies, together with Protein A/G beads or FLAG-M2 affinity gel and rotated at 4 °C for 2 h. For IP of endogenous proteins, FGFR1 and GPX4 were immunoprecipitated from lysates collected from three 10-cm dishes of HT1080 cells. Cells were lysed on ice, sonicated, and centrifuged at 4 °C for 10 min. An aliquot of the supernatant was reserved as the total protein fraction. The remaining lysates were incubated with the corresponding antibodies overnight at 4 °C. Protein A/G beads were then added to the lysate-antibody mixture and incubated for an additional 3 h at 4 °C. For both procedures, the beads were collected, washed three times with lysis buffer at 4 °C, resuspended in 2 × SDS sample buffer, boiled for 10 min, and subjected to immunoblotting analysis.

For immunoblotting, total cell lysates and immunoprecipitates were separated on 8-12% SDS-PAGE gels and transferred to polyvinylidene difluoride (PVDF) membranes (Millipore, Cat. IPVH00010). Membranes were blocked with 5% non-fat milk and incubated with specific primary antibodies (diluted in TBST supplemented with 5% BSA) overnight at 4 °C. After washing three times with TBST, membranes were incubated with horseradish peroxidase (HRP)-conjugated secondary antibodies for 1 h at room temperature with gentle agitation. Following three additional washes with TBST, protein bands were visualized using an enhanced chemiluminescence (ECL) detection system.

### RNA isolation and quantitative RT-PCR

Total RNA was extracted using TRIzol reagent (Thermo Fisher Scientific, Cat. 15596026CN) and reverse-transcribed to cDNA with the ReverTra Ace qPCR RT kit (TOYOBO, Cat. FSQ-101) according to the manufacturer’s instructions. Quantitative PCR was performed using the CFX96 Touch Real-Time PCR Detection System (Bio-Rad Laboratories, Inc., Hercules, USA) and the PerfectStart Green qPCR SuperMix (TransGen Biotech, Cat. AQ602). Relative gene expression levels were normalized to the housekeeping gene *Gapdh*. The primer sequences used are listed in **Supplementary Table S3**.

### Bimolecular fluorescence complementation (BiFC) assay

For BiFC assays, cells transfected with plasmids *GPX4*-VN155 and CC155-*FGFR1*, either individually or in combination, were plated on cover slides. Upon reaching 70% confluency, the cells were incubated at 4 °C for 2 h, washed with ice-cold PBS, and fixed with 4% paraformaldehyde for 30 min at 4 °C. The samples were mounted with Vectashield mounting medium, and fluorescent images were captured using a Nikon C2si confocal microscope.

### Proximity ligation assay (PLA)

HEK293T cells co-transfected with Flag-GPX4 and HA-FGFR1 were cultured on coverslips in 12-well cell culture plates. Cells were treated with vehicle or 5 nM FGF-1 for 15 min, followed by one wash with ice-cold PBS. After washing, cells were fixed with 4% (w/v) paraformaldehyde for 15 min at room temperature, then washed three additional times with PBS. Fixed cells were incubated with Duolink Blocking Solution (Sigma-Aldrich, Cat# DUO82007) for 90 min at 37 °C to block non-specific antibody binding. After blocking, cells were incubated overnight at 4 °C in a mixture of two primary antibodies: rabbit anti-HA and mouse anti-Flag. The PLA reaction was performed according to the manufacturer’s instruction. After developing PLA fluorescence signals, cell membranes were stained with the DIO membrane dye, and cell nuclei were counterstained with 4′,6-diamidino-2-phenylindole (DAPI). Fluorescence images were acquired on a Nikon C2si confocal laser scanning microscope.

### Plasmid construction

Full-length cDNAs encoding human FGFR1, FGFR2, FGFR3, FGFR4 and GPX4 were amplified by PCR using human cDNA templates from HEK293T cells. Site-directed mutagenesis of FGFR1 and GPX4 were performed using a PCR based method with PrimeSTAR HS polymerase (Takara, Cat. R010A). The resulting amplicons were cloned into the pcDNA3.3 vector for transfection or the pBOBI vector for lentivirus packaging. The lentiviral vector pLL3.7 was used to express shRNAs. The 19-nucleotide sequences for shRNA target sequences were as follows: human *FGFR1*, 5’-GTGGCTGTGAAGATGTTGA-3’ (1#) and 5’-GCATTGTGGAGAATGAGTA-3’ (2#); and mouse *Gpx4*, 5’-GCATCAAATGGAACTTTAA-3’. The pLL3.7-*Renilla* was used to express control shRNA (5’-GTAGCGCGGTGTATTATAC-3’). The plasmids containing fragments of YFP CDS (pBiFC-VN155I152L, Addgene, Cat. 27097 and pBiFC-CC155, Addgene, Cat. 22015) were kindly provided by Prof. Helen He Zhu (Shanghai Jiao Tong University School of Medicine).

For the BiFC experiment, VN155 was fused into the 3′end of the *GPX4* CDS sequence and CC155 was inserted into the *FGFR1* CDS between the signal peptide sequence and sequence encoding the extracellular domain. All constructs were verified by DNA sequencing. The primer sequences used for molecular cloning are listed in **Supplementary Table S4**.

### Expression and purification of recombinant human FGF-1, GPX4, and FGFRs

The cDNA encoding full-length wild-type human FGF-1 (FGF-1^WT^, residues 1-155) was subcloned into the pET-30a expression vector using NdeI and HindIII restriction sites. An N-terminal truncated FGF-1 variant (FGF-1^ΔNT^), lacking residues 1-24 (aa 25-155), was synthesized as a codon-optimized gene and cloned into the pET-30a vector via identical restriction enzyme sites by GenScript (Nanjing, China). FGF-1^WT^ and FGF-1^ΔNT^ were expressed in *Escherichia coli* strain BL21 (DE3). Transformed cells were grown at 37 °C until reaching an optical density at 600 nm (OD_600_) of 0.5. Protein expression was induced with 1 mM isopropyl β-D-1-thiogalactopyranoside (IPTG) for 4 h. Bacteria were harvested by centrifugation and lysed using an Emulsiflex-C3 high-pressure homogenizer (Avestin, Inc., Ottawa, Canada). The lysate was centrifuged to remove cell debris, and the soluble fractions containing FGF-1^WT^ or FGF-1^ΔNT^ were purified by sequentially heparin-affinity chromatography, followed by size-exclusion chromatography. Protein purity was assessed by SDS-PAGE and Coomassie Blue staining, and estimated to be >98% for all preparations.

Expression and purification of the His-tagged truncated human FGFR1 (aa 597-822), FGFR1 (aa 456-765), FGFR1-Y730F (aa 456-765) were performed as described^54^. Briefly, cDNAs for the truncated human FGFR1 were cloned into pET-28a or pET-Duet-1 vector, respectively, and transformed into *Escherichia coli* BL21 strain. The transformed *Escherichia coli* strain cells were cultured for 2 h at 37 °C with 1 mM IPTG to induce the expression of His-tagged truncated FGFR1. His-tagged proteins were purified by using Ni^2+^-NTA-agarose (Sigma-Aldrich, Cat. P6611).

The extracellular domains of human FGFR1 (residues 142-365), FGFR2 (residues 149-368), FGFR3 (residues 147-365), and FGFR4 (residues 144-355) were expressed in *Escherichia coli* BL21 (DE3) as inclusion bodies. Inclusion bodies were isolated and solubilized, and refolded *in vitro* through stepwise dialysis at 4 °C against a series of buffers: buffer A [25 mM Tris-HCl (pH 8.2), 150 mM NaCl, 7.5% glycerol], buffer B [25 mM Tris-HCl (pH 8.2), 100 mM NaCl, 5% glycerol], and buffer C [25 mM Tris-HCl (pH 8.2), 50 mM NaCl, 5% glycerol], with each step lasting at least 12 h. Refolded proteins were purified by heparin affinity chromatography followed by size-exclusion chromatography. All purification steps were performed out at 4 °C using an ÄKTA pure 25 L system (GE Healthcare).

His-tagged and GST-tagged human GPX4 constructs were generated with a Sec73Gly (U73G) mutation to enable recombinant expression in *Escherichia coli*, because selenocysteine incorporation requires specialized translation machinery that is not natively encoded in *E. coli*. The corresponding cDNAs were cloned into the pET-28a or pGEX-4T-1 vector, respectively, then the sequence-verified constructs were transformed into *Escherichia coli* BL21(DE3) competent cells. Recombinant protein expression was induced with 1 mM IPTG at 37 °C for 4 h. Following induction, cells were harvested and lysed, and inclusion bodies containing recombinant GPX4 were collected by centrifugation, then solubilized in 6 M guanidinium hydrochloride. GPX4 proteins were refolded by stepwise dialysis at 4 °C for 48 h: first against buffer A (25 mM HEPES, pH 8.0, 150 mM NaCl and 7.5% glycerol), then against buffer B (25 mM HEPES, pH 8.0, 100 mM NaCl and 5% glycerol). Refolded His-tagged GPX4 proteins were purified by Ni^2+^-affinity chromatography and further purified by size-exclusion chromatography. For GST-tagged GPX4, cells were harvested and lysed, and the soluble supernatant fraction was loaded onto glutathione sepharose beads (GE healthcare, Cat. 17513202) for affinity purification.

### GST pull-down assay

For the *in vitro* GST pull-down assay, bead-bound GST-GPX4 fusion proteins were incubated with purified His-tagged truncated FGFR1 (residues 597-822) in GST pull-down assay buffer [50 mM Tris-HCl (pH 7.6), 150 mM NaCl, 0.5% Nonidet P-40 (NP-40), and 1 mM PMSF] at 4 °C for 2 h. The beads were washed three times with the same buffer and resuspended in SDS loading buffer for immunoblotting.

### Surface plasmon resonance (SPR) spectroscopy

SPR measurements were performed on a Biacore T200 instrument (GE Healthcare) in HBS-EP buffer [10 mM HEPES (pH 7.4), 150 mM NaCl, 3 mM EDTA, 0.005% P20]. For kinase interaction studies, the phosphorylated human FGFR1 kinase domain (p-FGFR1, p-FGFR1-Y653/654 and p-FGFR1-Y730F, residues 455-765) was immobilized on a CM5 sensor chip (Cytiva, Cat. BR100530) by amine coupling to a density of 300-400 response units (RU). A reference flow cell was prepared simultaneously. Full-length GPX4 was injected at increasing concentrations (5-200 μg/mL) using a multi-cycle kinetic protocol (association 180 s, dissociation 180 s) at 50 μL/min.

For ligand-receptor binding analysis, the extracellular domains of human FGFR1 (residues 142-365), FGFR2 (residues 149-368), FGFR3 (residues 147-365) and FGFR4 (residues 144-355) were individually immobilized on CM5 sensor chips. Recombinant human FGF-1^WT^ and FGF-1^ΔNT^ were injected at increasing concentrations using a multi-cycle kinetic protocol (association 180 s, dissociation 180 s) at a flow rate of 50 μL/min. The sensor chip surfaces were regenerated with high-salt buffers between injections. Data were processed with BIAEvaluation software, and equilibrium dissociation constants (Kd) were calculated from fitted saturation binding curves.

### Mass spectrometry analysis

To identify proteins that interact with FGFR1, Flag-tagged FGFR1 was expressed in cardiomyocytes from *Fgfr1*-CKO mice. Following immunoprecipitation with an anti-Flag antibody, proteins were separated by SDS-PAGE and visualized using Coomassie blue staining. The stained gels were excised and digested by trypsin, followed by mass spectrometry analysis.

### *In vitro* kinase assay

HA-tagged WT-or kinase-dead forms of FGFR1 (KD-FGFR1, FGFR1-Y653/654F) were overexpressed in HEK293T cells and immunoprecipitated with HA magnetic beads (Santa Cruz Biotechnology, Cat. sc-500773) in lysis buffer. The purified HA-tagged WT-FGFR1 or KD-FGFR1 proteins were then incubated with bacterially expressed His-tagged GPX4 or/and 2YF-GPX4 in a kinase reaction buffer [25 mM HEPES (pH 7.4), 50 mM NaCl, 5 mM MgCl_2_, 1 mM DTT, 0.5 mg/ml BSA, 1 mM Na_3_VO_4_] supplemented with 100 μM cold ATP (Sigma-Aldrich, Cat. A7699) and 0.5 μCi ^32^P-labeled ATP (PerkinElmer, Cat. NEG502A250UC) for 30 min at 28 °C. The kinase assay was terminated by the addition of SDS sample buffer and analyzed by immunoblotting.

### GPX4 activity assay

To measure the activity of ectopically expressed GPX4, we used full-length Flag-tagged WT-, 2YF-, 2YD-, or 2YE-GPX4 constructs (nucleotides 1-801). All constructs retained the native selenocysteine insertion sequence (SECIS) element in the 3’ untranslated region (UTR) to enable selenocysteine incorporation at residue 73. These constructs were individually co-infected with or without FGFR1 into HEK293T or *GPX4*^−/−^HT1080 cells. Cells were harvested 24 h after transfection. Flag-or HA-tagged GPX4 was immunoprecipitated using Flag antibody-conjugated beads for 3 h at 4 °C. The immunoprecipitates were washed with GPX4 assay buffer [100 mM Tris-HCl (pH 8.0), 2 mM EDTA, 0.1% peroxide-free Triton-X100] three times. The Flag-or HA-tagged proteins were eluted from beads with Flag or HA peptide in elution buffer [25 mM Tris-HCl (pH 8.0), 150 mM NaCl] supplemented with a protease inhibitor cocktail. Tert-butyl hydroperoxide (TBHP) was used as the substrate for GPX4 activity detection.

To measure endogenous GPX4 activity, HT1080 cells treated with RSL3 with or without FGFR1 inhibitor (PD166866 or PD173074) or WT and *FGFR1*^−/−^HT1080 cells reconstituted with empty vector or HA-FGFR1 were cultured in 10-cm dishes. Following treatment with the indicated drugs, cells were harvested via trypsinization, washed twice with ice-cold PBS, and cell pellets were lysed in GPX4 assay buffer. Lysate supernatants were collected for total protein quantification, and 1-palmitoyl-2-hydroperoxyoctadecadienoyl-sn-glycero-3-phosphocholine (PCOOH) was used as the specific substrate for endogenous GPX4 activity detection.

GPX4 activity was determined using a coupled enzymatic assay by monitoring the consumption of NADPH, as described^14,56^. Briefly, cells lysates were sonicated and centrifuged at 12,000 rpm for 10 min at 4 °C to remove debris. Supernatant total protein concentration was measured with a standard protein assay. A total reaction volume of 200 µL, containing protein-normalized lysates or eluted proteins, was prepared in a 96-well plate. The reaction mixture was prepared by adding reduced glutathione (GSH) (5 mM), glutathione reductase (GR) (1.5 U/mL) and NADPH (0.25 mM). Enzymatic reactions were initiated by adding TBHP (3 mM or the indicated concentration) or PCOOH (50 μM). The rate of NADPH absorbance decline was measured kinetically at 340 nm, with reads collected every 10 s for 10 min using a microplate reader. GPX4 activity was calculated from the slope of NADPH oxidation.

### Immunohistochemistry

For immunohistochemistry, 5 μm-thick sections were baked at 75 °C for 4 h. The sections were then deparaffinized and rehydrated using xylene and a graded ethanol series. After washing with PBS, antigen retrieval was performed by boiling the sections in 10 mM sodium citrate buffer (pH 6.0) at 100 °C for 30 min. The sections were rinsed twice with PBS and treated with 3% H_2_O_2_ for 30 min to block endogenous peroxidase activity, and blocked with 1% BSA for 1 h at room temperature, followed by incubation with 4-HNE or Col1α1 antibodies overnight at 4 °C. Following washes with PBS, sections were incubated with HRP-conjugated goat anti-rabbit or mouse secondary antibodies for 1 h at room temperature. Signal detection was performed using the DAB Peroxidase (HRP) Substrate Kit (Proteintech, Cat. PR30018), and then sections were counterstained with hematoxylin according to the manufacturer’s protocol.

### (Oxy)Lipidomics analysis

Lipids were extracted from mouse heart tissues using a modified methyl-*tert*-butyl ether (MTBE) method^57^. Briefly, 25 mg ± 1 mg (wet weight) frozen heart tissues were combined with homogenization beads with 200 μL ultrapure water, then vortexed for 30 s. The sample was homogenized at 35 Hz for 4 min, followed by ultrasonication in an ice-water bath at 4 °C for 5 min; this homogenization-sonication cycle was repeated three times. Next, 480 μL of extraction solvent (5:1, v/v, MTBE: methanol) containing deuterated lipid internal standards was added to the homogenate. The mixture was vortexed for 30 s, ultrasonicated at 4 °C for 10 min, and then incubated at −40 °C for 1 h to precipitate protein debris. After incubation, the sample was centrifuged at 900 *g* for 15 min at 4 °C. The supernatant was collected, evaporated to dryness under vacuum, and reconstituted in 200 μL of dichloromethane: methanol (1:1, v/v). The reconstituted extract was vortexed for 30 s, ultrasonicated at 4 °C for 10 min, and centrifuged at 13,800 *g* for 15 min at 4 °C to remove insoluble debris. A 75 μL aliquot of the final supernatant was transferred to glass vials for instrumental analysis. Quality control (QC) samples were prepared by pooling equal volumes of supernatant from all individual experimental samples.

LC-MS/MS analysis was performed on a Vanquish ultra-high-performance liquid chromatography (UHPLC) system (Thermo Fisher Scientific) coupled to an Orbitrap Exploris 120 mass spectrometer equipped with an electrospray ionization (ESI) source. Lipids separation was performed on a Phenomenex Kinetex C18 column (2.1 mm × 100 mm, 2.6 μm). Mobile phase A consisted of water: acetonitrile (6:4, v/v) with 10 mM ammonium formate, and mobile phase B consisted of isopropanol: acetonitrile (9:1, v/v) with 10 mM ammonium formate. The injection volume was set to2 μL, and data-dependent acquisition was controlled by Xcalibur software (Thermo Fisher Scientific). ESI source parameters were set as follows: sheath gas = 30 Arb, auxiliary gas = 10 Arb, capillary temperature = 320 °C, full MS resolution = 60,000, MS/MS resolution = 15,000, stepped normalized collision energy (SNCE) = 15/30/45, spray voltage = 3.8 kV (positive ion mode) and −3.4 kV (negative ion mode). Raw MS data files were converted to mzXML format using msconvert (ProteoWizard). Peak picking, extraction, alignment and integration were performed using the CentWave algorithm implemented in the XCMS package in R. Annotation parameters were set as minfrac = 0.5 and cutoff = 0.3. Lipid identification was performed by spectral matching against the LipidBlast library. All downstream analyses were restricted to annotated oxidized lipid species.

After noise filtering based on relative standard deviation (RSD), missing peak intensities for oxidized lipids were imputed as half the minimum detected intensity for each oxidized lipid feature. Total ion current (TIC) normalization was applied prior to statistical analysis. The processed dataset, containing peak identifiers, sample labels, and normalized peak areas for oxidized lipids, was imported into the R package ropls (v1.34.0). Data were log-transformed and auto-scaled to reduce the influence of technical noise and skewed feature intensity distributions. Unsupervised principal component analysis (PCA) was used to visualize sample clustering based on oxidized lipid profiles, and potential outliers were screened using a 95% confidence interval. Supervised orthogonal projections to latent structures-discriminant analysis (OPLS-DA) was conducted to characterize intergroup separation and identify differentially abundant oxidized lipid species. Model performance was evaluated via seven-fold cross-validation, which calculated R^2^ (explained variance) and Q^2^ (predicted variance) values; 200-fold permutation testing was carried out to assess model robustness and avoid overfitting. Oxidized lipids with variable importance in projection (VIP) > 1 and *p* < 0.05 (two-tailed Student’s *t* test) were defined as significantly altered.

### Statistical analyses

For experiments with only two groups, Student’s *t* test was used for statistical comparisons. Pearson’s correlation test was used to measure the strength of association between two variables. ANOVA with Tukey’s post-test (one-way ANOVA for comparisons between groups, two-way ANOVA for comparisons of magnitude of changes between different groups from different cell lines or treatments) was used to compare values among different experimental groups using the GraphPad Prism 10.

## Data availability statement

Source data for the Figures and Extended Data Figures are provided with the paper. Further data supporting the findings of this study are available from the corresponding author upon reasonable request.

## Acknowledgements

This work was supported by grants from the National Natural Science Foundation of China (82522089 and 82373934 to L.T.S., 82530107, U25A2036 and 92357304 to Z.F.H., U25D9022 and U22A20385 to X.K.L., 82504890 to L.Y.W.), the Innovative Drug Research and Development National Science and Technology Major Project (2025ZD1802000 to Z.F.H., 2026ZD1805501 to L.T.S.), and Postdoctoral Science Preferential Funding of Zhejiang Province (ZJ2025238 to L.Y.W.). We would like to express our sincere gratitude to Professor Fudi Wang and Junxia Min (Zhejiang University) for their constructive suggestions and insightful comments on this manuscript. We also thank the Scientific Research Center of Wenzhou Medical University for consultation and instrument availability that supported this work.

## Author contributions

L.T.S., Z.F.H. and X.K.L. conceived the project and designed the experiments. L.Y.W., W.L.D., J.Q., S.S.Z., H.X.L., Y.S., S.Y.X., Y.S.H., S.L.T., J.C., H.Y., J.Z.X., W.N.L. and L.F. performed most of the experiments and participated in discussions regarding the results. J.Q. and S.L.T. expressed and purified proteins and performed SPR. L.T.S., Z.F.H. and L.Y.W. analyzed data and wrote the paper.

## Additional information

Supplementary information is available for this paper.

Correspondence and requests for materials should be addressed to Z.F. H.

---

**Extended Data Fig. 1.**
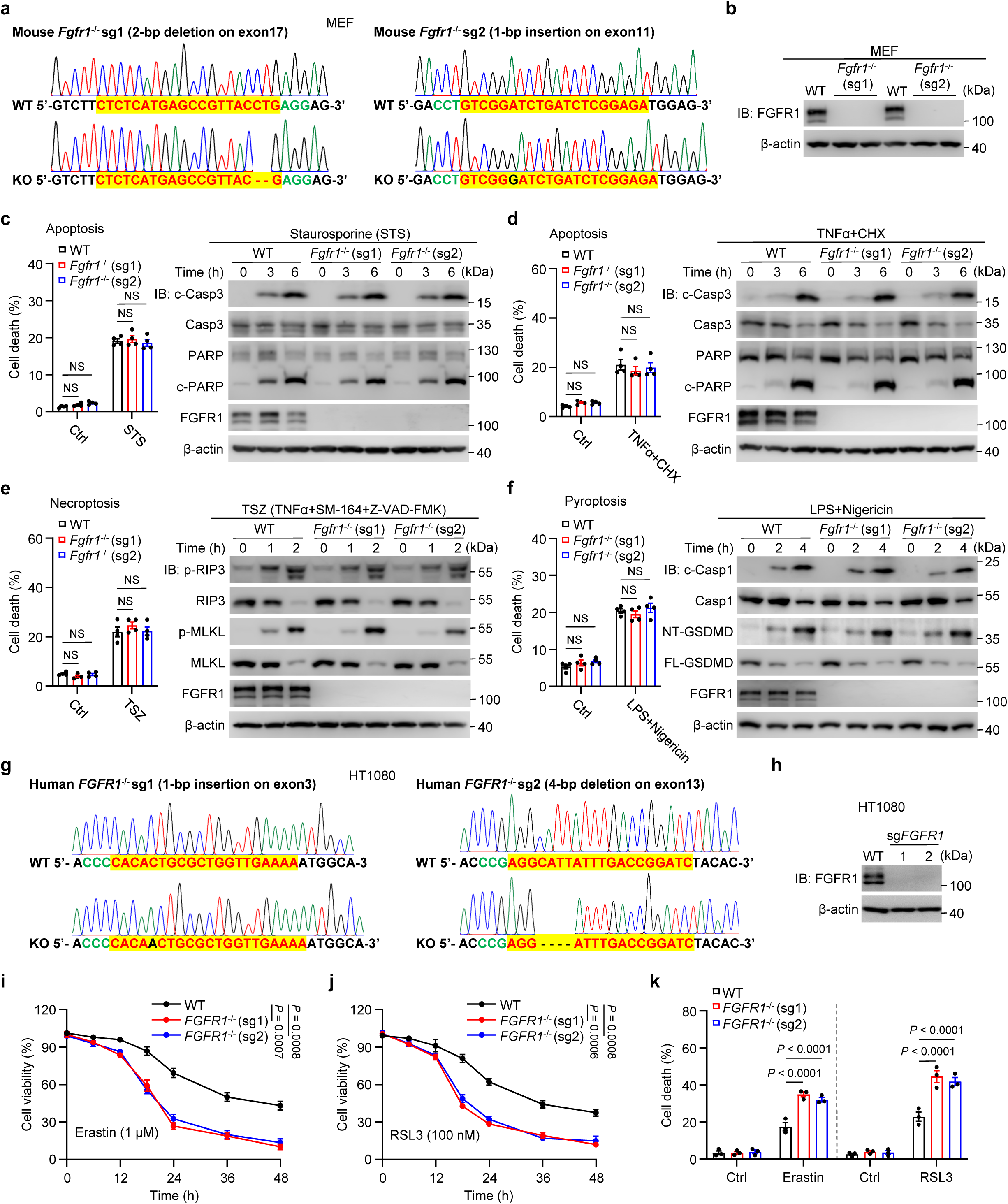
Deficiency of *Fgfr1* promotes ferroptosis. **a,** Schematic of CRISPR-Cas9-mediated knockout (KO) of *Fgfr1* (*Fgfr1*^−/−^) in mouse embryonic fibroblasts (MEFs). Two distinct sets of sgRNAs (sg1 or sg2) were applied to generate *Fgfr1*^−/−^MEFs. The gRNA-targeting sequences used are highlighted in yellow, and the protospacer-adjacent motif (PAM) sequence is indicated in green. **b,** Immunoblot analysis of FGFR1 levels in CRISPR control (WT) or *Fgfr1*^−/−^ MEFs. **c-f,** Cell death (left) and immunoblot analysis (right) of WT and *Fgfr1*^−/−^ (sg1 and sg2) MEFs treated with 0.5 μM staurosporine (STS, **c**), 1 μg/ml cycloheximide (CHX) plus 10 ng/ml TNFα (**d**), 10ng/ml TNFα in combination with 0.1 μM SM-164 and 20 μM Z-VAD-FMK (TSZ, **e**), or 1 μg/ml LPS together with 20 μM nigericin (**f**) for the indicated durations. **g,** Schematic of CRISPR-Cas9-mediated KO of *FGFR1* (*FGFR1*^−/−^) in HT1080 cells. **h,** Immunoblot analysis of FGFR1 levels in WT or *FGFR1*^−/−^HT1080 cells. **i,j,** Cell viability of WT or *FGFR1*^−/−^ HT1080 cells subjected to the indicated treatments. **k,** Cell death were assessed in WT and *FGFR1*^−/−^ HT1080 cells treated with 10 μM Erastin or 0.5 μM RSL3. *n* = 3-4 biologically independent experiments in (left graphs of **c-f, i-k**). Data are mean ± s.e.m.; ordinary two-way ANOVA, followed by Sidak in (left graphs of **c-f, k**); two-way ANOVA (repeated measure) followed by Tukey in (**i, j**); \*\*\**p* < 0.001, \*\*\*\**p* < 0.0001, NS, not significant; Gel source data blots can be found in Supplementary Fig. 1.

**Extended Data Fig. 2.**
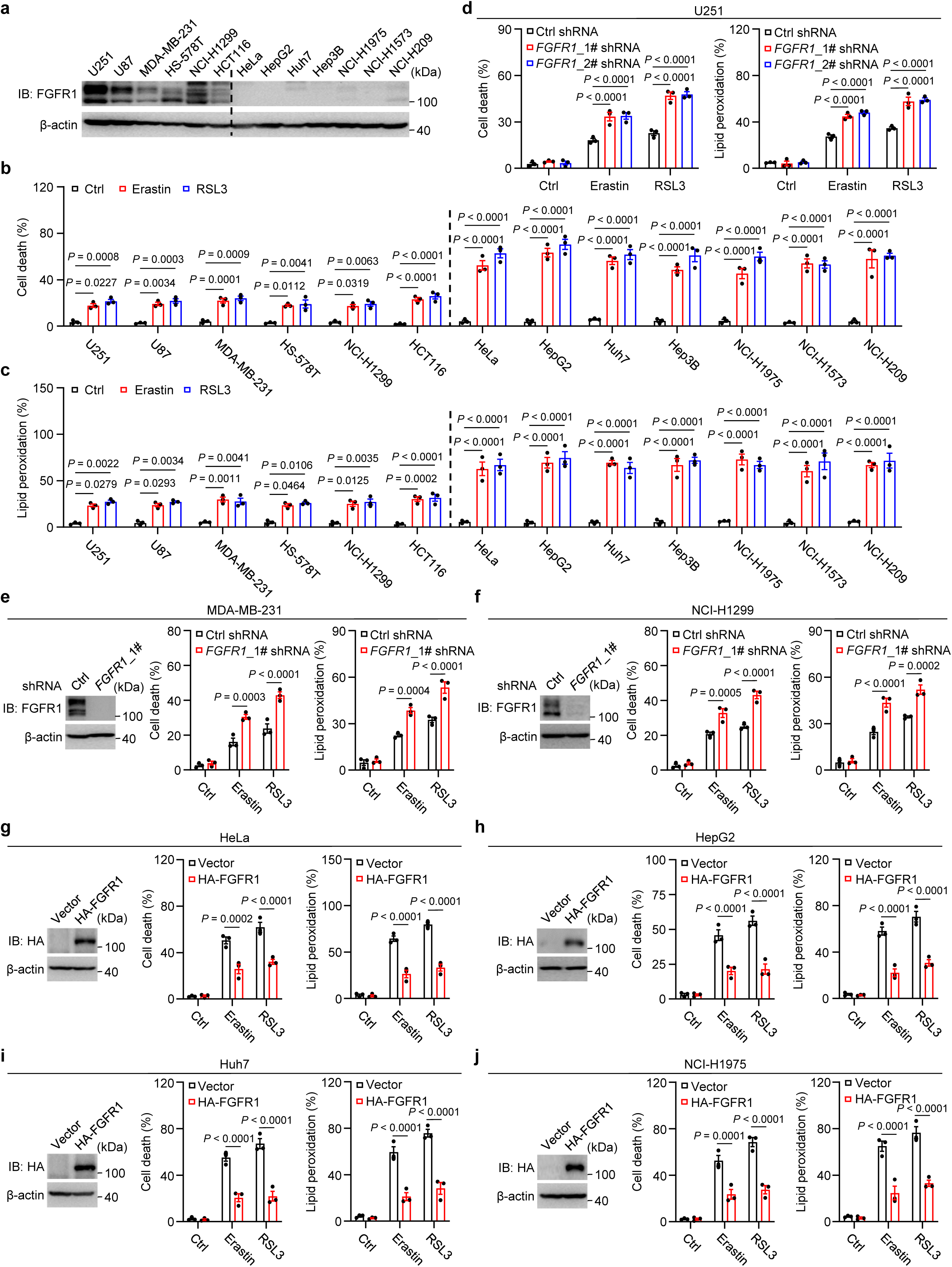
FGFR1 has a direct impact on ferroptosis sensitivity. **a,** The protein levels of FGFR1 in the indicated cell lines. **b,c,** Cell death (**b**) and lipid peroxidation (**c**) measurements in the indicated cell lines treated with Erastin or RSL3. **d-f,** Knockdown of *FGFR1* exacerbates cell death and lipid peroxidation in U251 (**d**), MDA-MB-231 (**e**) and NCI-H1299 (**f**) cells subjected to the indicated treatments. **g-j,** Overexpression of HA tagged FGFR1 (HA-FGFR1) suppresses the cell death and lipid peroxidation in HeLa (**g**), HepG2 (**h**) Huh7 (**i**), and NCI-H1975 (**j**) cells. For panels **e-j**, immunoblots (left) show FGFR1 levels in the indicated cells. Graphs show cell death (middle) and lipid peroxidation (right) measurements for the indicated cells treated with 0.5 μM RSL3 or 10 μM Erastin. *n* = 3 biologically independent experiments in (**b-j**). Data are mean ± s.e.m.; ordinary two-way ANOVA, followed by Sidak in (**b-d**, middle and right graphs of **e-j**); \**p* < 0.05, \*\**p* < 0.01, \*\*\**p* < 0.001, \*\*\*\**p* < 0.0001; Gel source data blots can be found in Supplementary Fig. 1.

**Extended Data Fig. 3.**
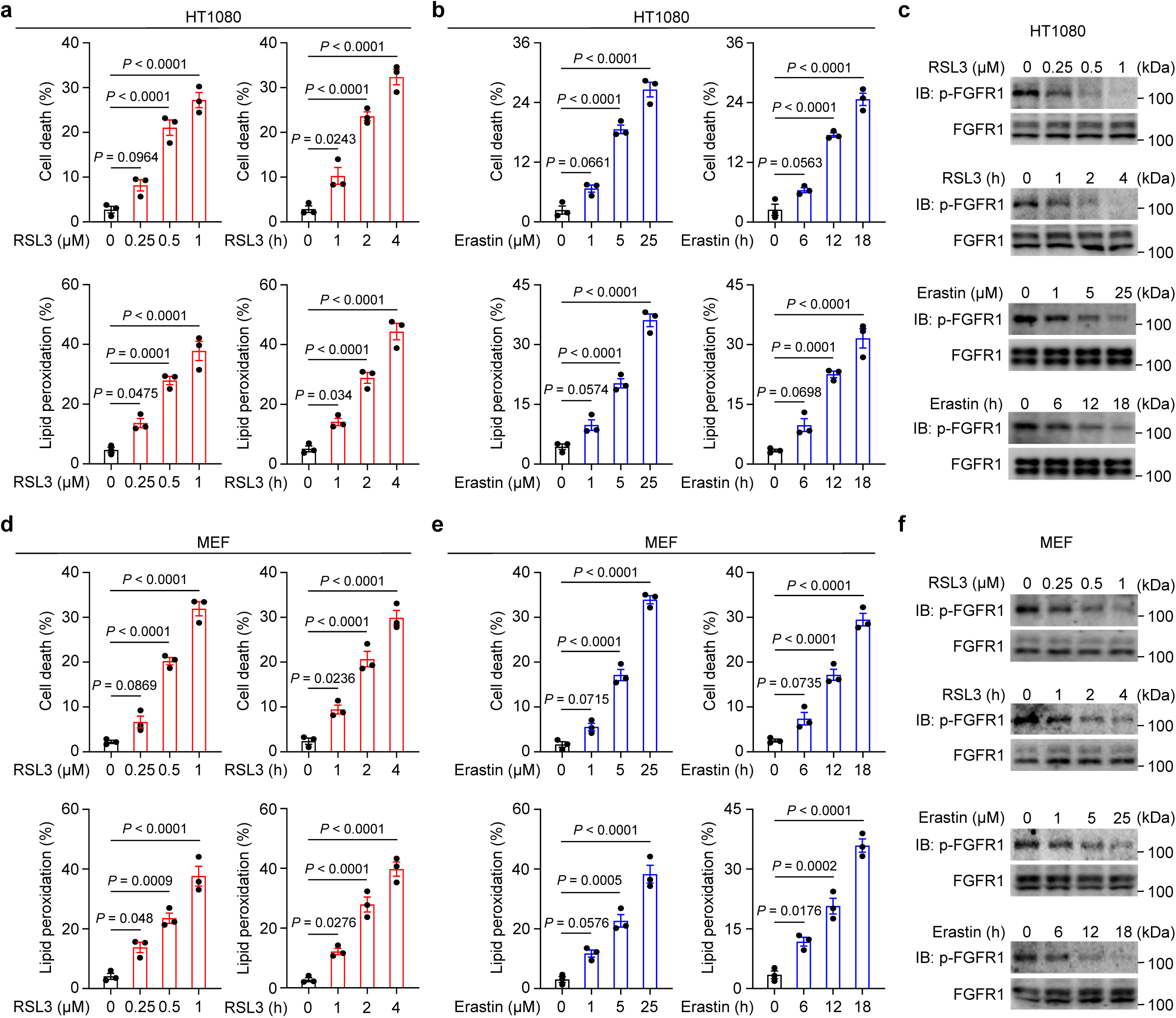
Ferroptosis suppresses FGFR1 kinase activity. **a,b,d,e,** Dose-and time-dependent cell death and lipid peroxidation in HT1080 (**a, b**) and MEF (**d, e**) cells treated with RSL3 (**a, d**) or Erastin (**b, e**). **c,f,** Immunoblot analysis of total cell lysates from HT1080 (**c**) or MEF (**f**) cells treated with RSL3 or Erastin for the indicated durations. *n* = 3 biologically independent experiments in (**a, b, d, e**). Data are mean ± s.e.m.; ordinary one-way ANOVA, followed by Tukey in (**a, b, d, e**); \**p* < 0.05, \*\**p* < 0.01, \*\*\**p* < 0.001, \*\*\*\**p* < 0.0001; Gel source data blots can be found in Supplementary Fig. 1.

**Extended Data Fig. 4.**
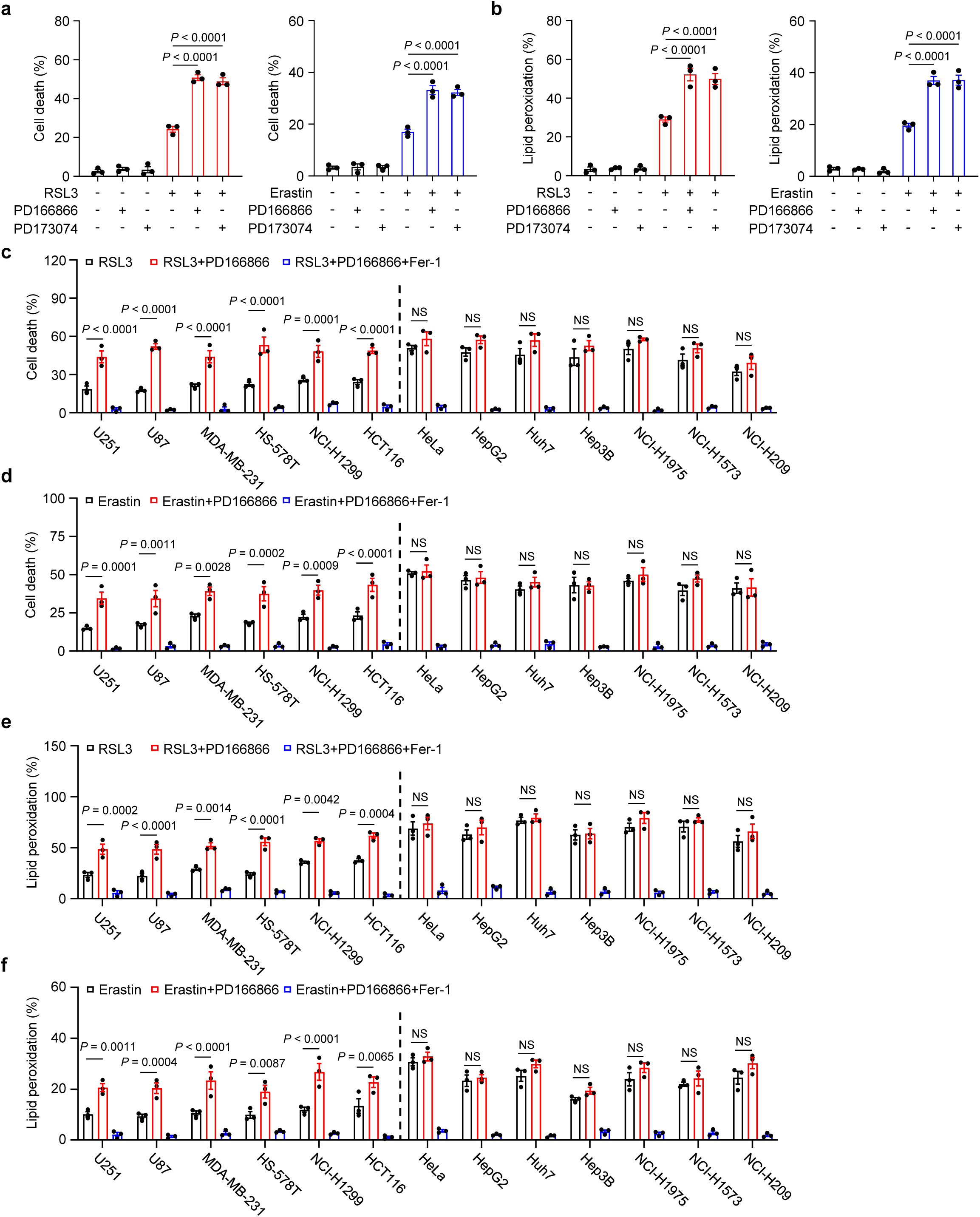
Inhibition of FGFR1 kinase activity promotes ferroptosis. **a,b,** The measurement of cell death (**a**) and lipid peroxidation (**b**) in HT1080 cells treated with 0.5 μM RSL3 or 10 μM Erastin following pretreatment with FGFR1 inhibitors (PD166866 and PD173074). **c-f,** Cell death (**c, d**) and lipid peroxidation (**e, f**) measurements for the indicated cell lines treated with 0.5 μM RSL3 (**c, e**) or 10 μM Erastin (**d, f**) with or without PD166866 or Fer-1. *n* = 3 biologically independent experiments in (**a-f**). Data are mean ± s.e.m.; ordinary one-way ANOVA, followed by Tukey in (**a, b**); ordinary two-way ANOVA, followed by Sidak in (**c-f**); \*\**p* < 0.01, \*\*\**p* < 0.001, \*\*\*\**p* < 0.0001, NS, not significant.

**Extended Data Fig. 5.**
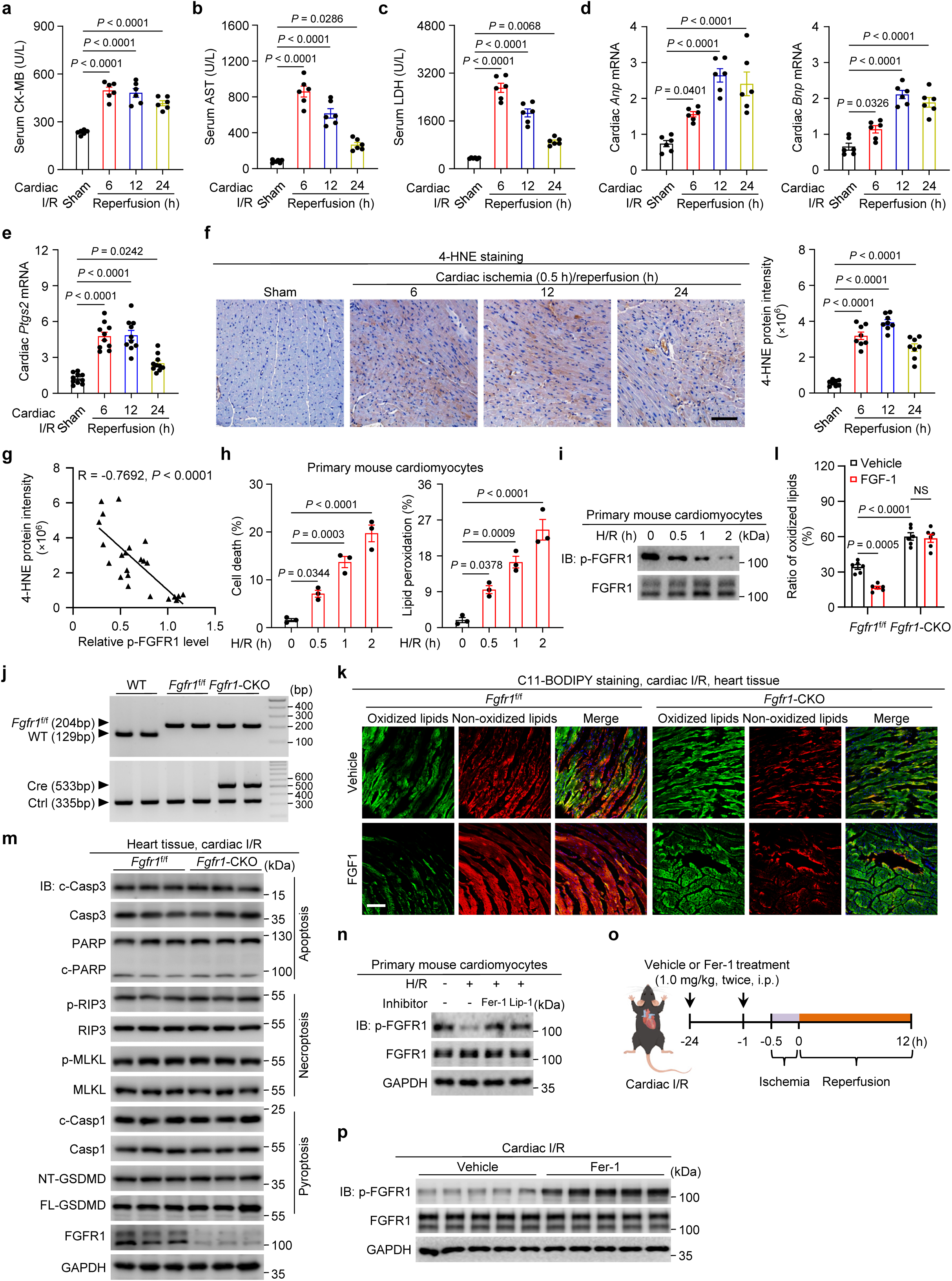
Cardiac ischemia-reperfusion (I/R) induced ferroptosis. **a-c,** Serum creatine kinase-MB isoenzyme (CK-MB, **a**), aspartate transaminase (AST, **b**), and lactate dehydrogenase (LDH, **c**) levels in C57BL/6J mice subjected to sham surgery or 0.5 h/6, 12, or 24 h cardiac I/R injury. *n* = 6 mice/group. **d,e,** Relative mRNA levels of *Anp* and *Bnp* (**d**, *n* = 6 mice/group), or *Ptgs2* (**e**, *n* = 10 mice/group) in the indicated mice. Relative mRNA levels were normalized to *Gapdh*. **f,** Representative images and quantitative analyses of cardiac sections stained with 4-HNE in the indicated mice subjected to cardiac I/R. *n* = 8 mice/group. Scale bar, 100 μm. **g,** Correlation between 4-HNE protein levels and phosphorylated FGFR1 (Tyr653/654, p-FGFR1) in cardiac tissues from the indicated mice. Ratios of p-FGFR1 to total FGFR1 was calculated. *n* = 24 mice. **h,** Cell death (left) and lipid peroxidation (right) in primary mouse cardiomyocytes from C57BL/6J mice subjected to hypoxia/reoxygenation (H/R) challenge. *n* = 3 biologically independent experiments. **i,** Immunoblot analysis of total cell lysates from primary mouse cardiomyocytes upon H/R challenge. **j,** PCR genotyping confirming successful cardiac-specific *Fgfr1* knockout in mice. **k,l,** Representative images (**k**) and quantification (**l**) of C11-BODIPY staining in heart sections from the indicated groups. Scale bar, 100 μm. *n* = 6 mice/group. **m,** Immunoblot analysis of apoptosis, necroptosis, pyroptosis-related proteins in heart tissues from *Fgfr1*^f/f^ and *Fgfr1*-CKO mice subjected to cardiac I/R. **n,** Immunoblot analysis of p-FGFR1 levels in primary mouse cardiomyocytes from C57BL/6J mice subjected to H/R challenge in the presence of ferroptosis inhibitors Fer-1 or Lip-1. **o,** Schematic of the Fer-1 treatment regimen in C57BL/6 mice. Mice received Fer-1 or saline at the 24 h and 0.5 h before ischemia, followed by cardiac I/R. **p,** Immunoblot analysis of total cardiac lysates from Fer-1-treated C57BL/6J mice after cardiac I/R treatment. Data are mean ± s.e.m.; ordinary one-way ANOVA, followed by Tukey in (**a-e, h,** right graph of **f**); Pearson’s correlation in (**g**); ordinary two-way ANOVA, followed by Tukey in (**l**); \**p* < 0.05, \*\**p* < 0.01, \*\*\**p* < 0.001, \*\*\*\**p* < 0.0001, NS, not significant; Gel source data blots can be found in Supplementary Fig. 1.

**Extended Data Fig. 6.**
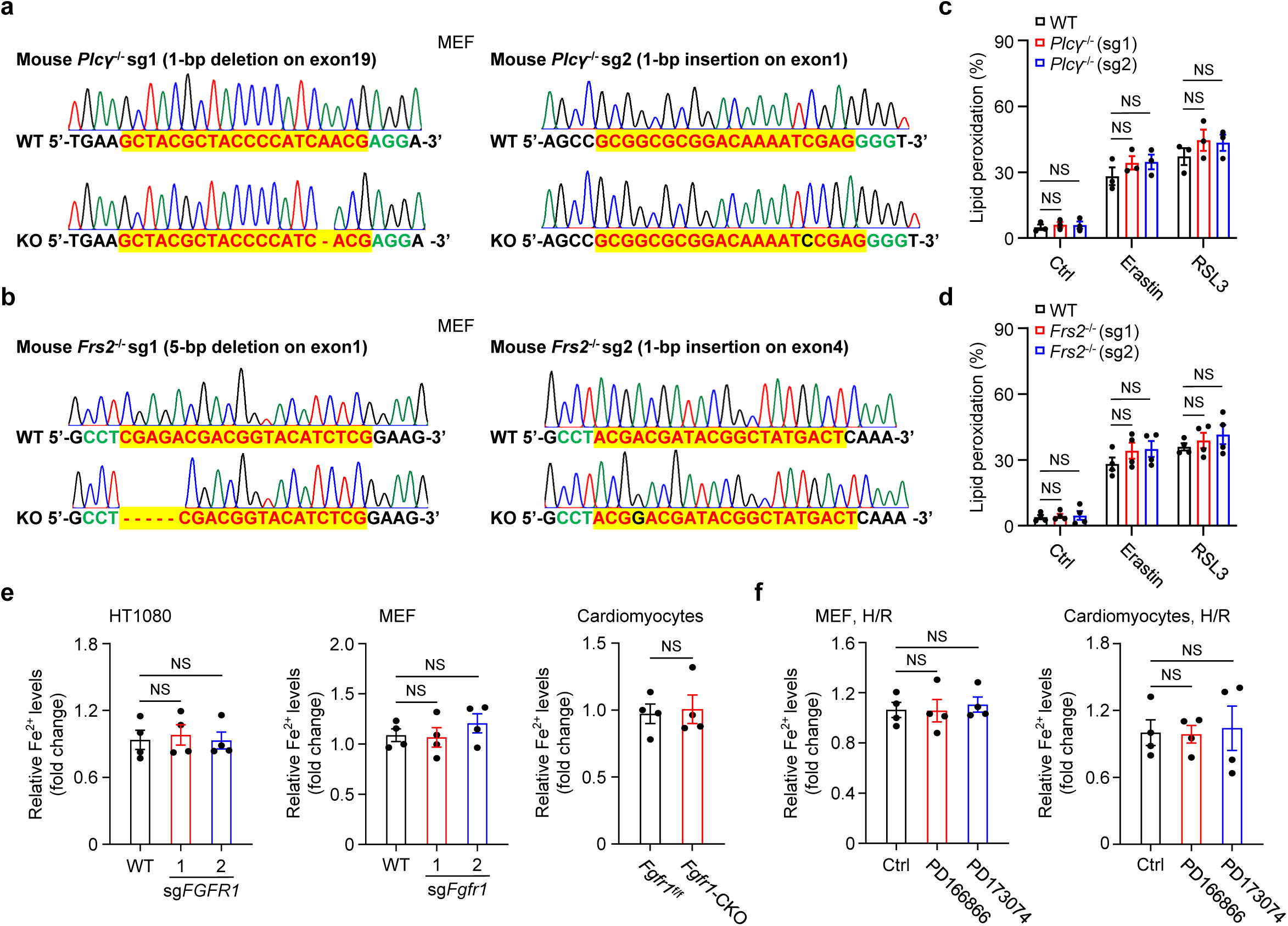
Deficiency of *Plcγ or Frs2* does not affect ferroptosis. **a,b,** Schematic of CRISPR-Cas9-mediated knockout of *Plcγ* (*Plcγ*^−/−^, **a**) or *Frs2* (*Frs2*^−/−^, **b**) in MEFs. Two distinct sets of sgRNAs were applied to generate *Plcγ*^−/−^or *Frs2*^−/−^MEFs. The gRNA-targeting sequences used are highlighted in yellow and the PAM sequence is indicated in green. **c,d,** Lipid peroxidation in WT and *Plcγ*^−/−^ (**c**) or *Frs2*^−/−^ (**d**) MEFs treated with 10 μM Erastin or 0.5 μM RSL3. **e,** The levels of cellular divalent iron in the WT and *Fgfr1*^−/−^ HT1080 cells, MEFs, or cardiomyocytes. **f,** Cellular divalent iron levels in MEFs or cardiomyocytes treated with FGFR1 inhibitors upon H/R treatment. *n* = 4 biologically independent experiments in (**c-f**). Data are mean ± s.e.m.; ordinary two-way ANOVA, followed by Sidak in (**c, d**); ordinary one-way ANOVA, followed by Tukey in (left and middle of graphs of **e, f**); Two-tailed unpaired Student *t* test (right graph of **e**); NS, not significant.

**Extended Data Fig. 7.**
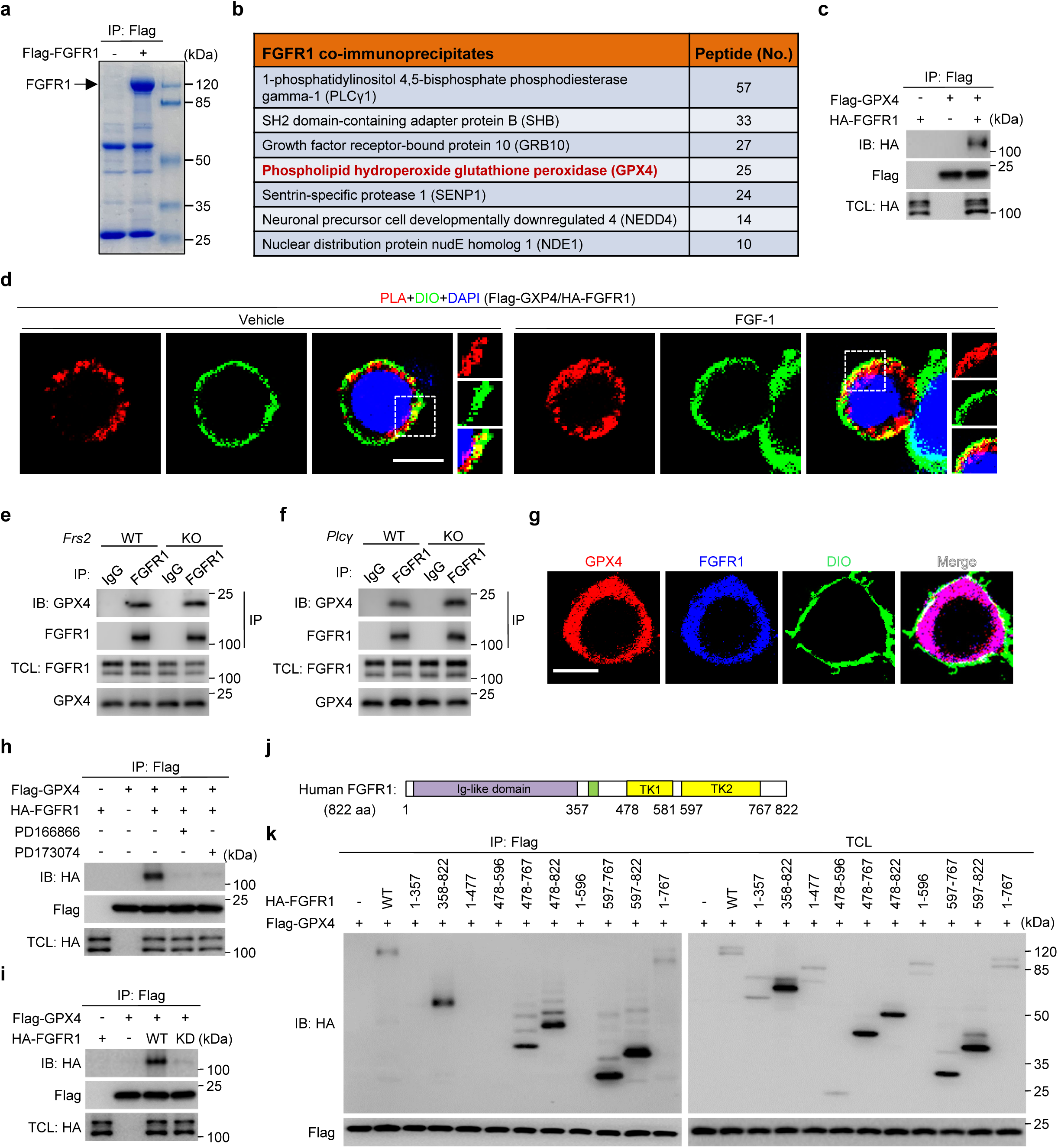
FGFR1 directly interacts with GPX4. **a,** Coomassie blue staining of FGFR1-interacting proteins. Flag-tagged FGFR1 was expressed in the cardiomyocytes from *Fgfr1*-CKO mice. **b,** A list of FGFR1-associated proteins identified by mass spectrometry (MS). **c,** Interaction between ectopically expressed Flag-GPX4 and HA-FGFR1. HEK293T cells were lysed and subjected to immunoprecipitation (IP) against Flag, followed by immunoblotting. TCL, total cell lysate. **d,** PLA analysis of the interaction between FGFR1 and GPX4 in the cell membrane. The cells were transfected with Flag-GPX4 and HA-FGFR1, followed by FGF-1 treatment and then subjected to PLA analysis. DIO was used as a cell membrane marker. Scale bars, 10 μm. **e,f,** Interaction between endogenous FGFR1 and GPX4. FGFR1 in WT and *Plcγ*^−/−^ (**e**) or *Frs2*^−/−^ (**f**) MEFs were immunoprecipitated (IP), followed by immunoblotting with antibodies indicated. **g,** Co-localization of GPX4 and FGFR1 on the cell membrane of HT1080 cells. The HT1080 cells were stained for FGFR1, GPX4 and DIO. Scale bars, 10 μm. **h,** FGFR1 inhibitors (PD166866 and PD173074) impaired the association between FGFR1 and GPX4. **i,** FGFR1 kinase activity is required for its interaction with GPX4. HA-tagged WT-FGFR1 or KD-FGFR1 was co-expressed with Flag-GPX4 in HEK293T cells. **j,** Diagram of human FGFR1. TK, tyrosine kinase domain. **k,** The TK2 domain (aa 597-767) of FGFR1 is mainly responsible for its interaction with GPX4. Gel source data blots can be found in Supplementary Fig. 1.

**Extended Data Fig. 8.**
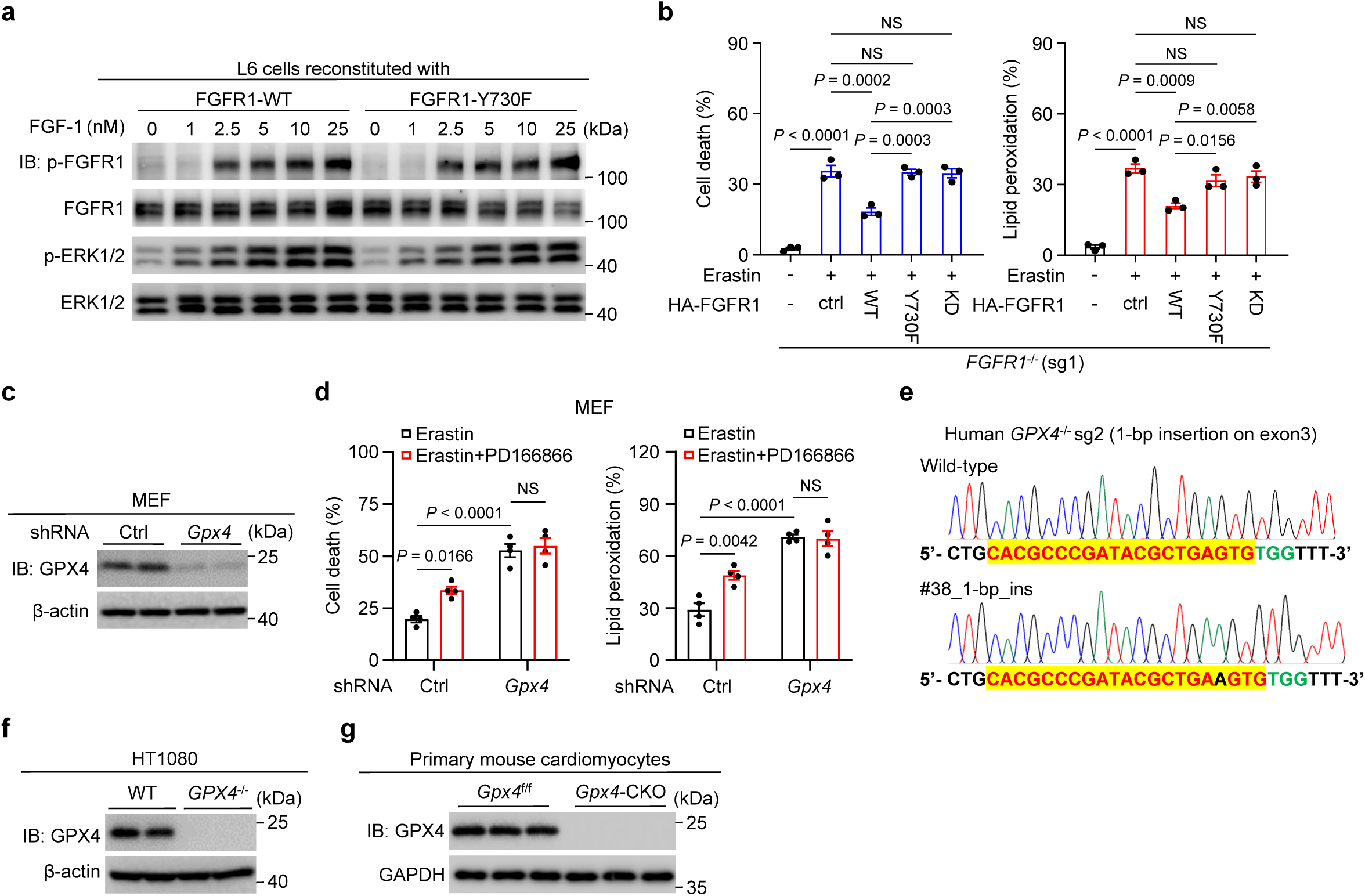
FGFR1 modulates ferroptosis via GPX4. **a,** Immunoblots showing dose-dependent activation of FGF/FGFR signaling pathway by FGF-1 in the L6 cells stably expressing FGFR1-WT or FGFR1-Y730F. **b,** Cell death and lipid peroxidation were performed in *FGFR1*^−/−^ HT1080 cells reconstituted with empty vector (ctrl), and HA-tagged WT-FGFR1, Y730F-FGFR1 or KD-FGFR1 subjected to 10 μM Erastin treatments. **c,** Knockdown of *Gpx4* in MEF cells. **d,** Measurement of cell death (left) or lipid peroxidation (right) in WT and *Gpx4*-KD MEFs treated with Erastin with/without the FGFR1 inhibitor PD166866. **e,** Generation of *GPX4*-knockout (*GPX4*^−/−^) HT1080 cell line. Sequencing results showing the 1-bp insertion found in clone #38. **f,** Immunoblot analysis of total proteins from CRISPR control (WT) or *GPX4*^−/−^ HT1080 cells. **g,** Immunoblot analysis of GPX4 in primary mouse cardiomyocytes isolated from *Gpx4*^f/f^ and *Gpx4*-CKO mice. *n* = 3-4 biologically independent experiments in (**b, d**). Data are mean ± s.e.m.; ordinary one-way ANOVA, followed by Tukey in (**b**); ordinary two-way ANOVA, followed by Tukey in (**d**). \**p* < 0.05, \*\**p* < 0.01, \*\*\**p* < 0.001, \*\*\*\**p* < 0.0001, NS, not significant; Gel source data blots can be found in Supplementary Fig. 1.

**Extended Data Fig. 9.**
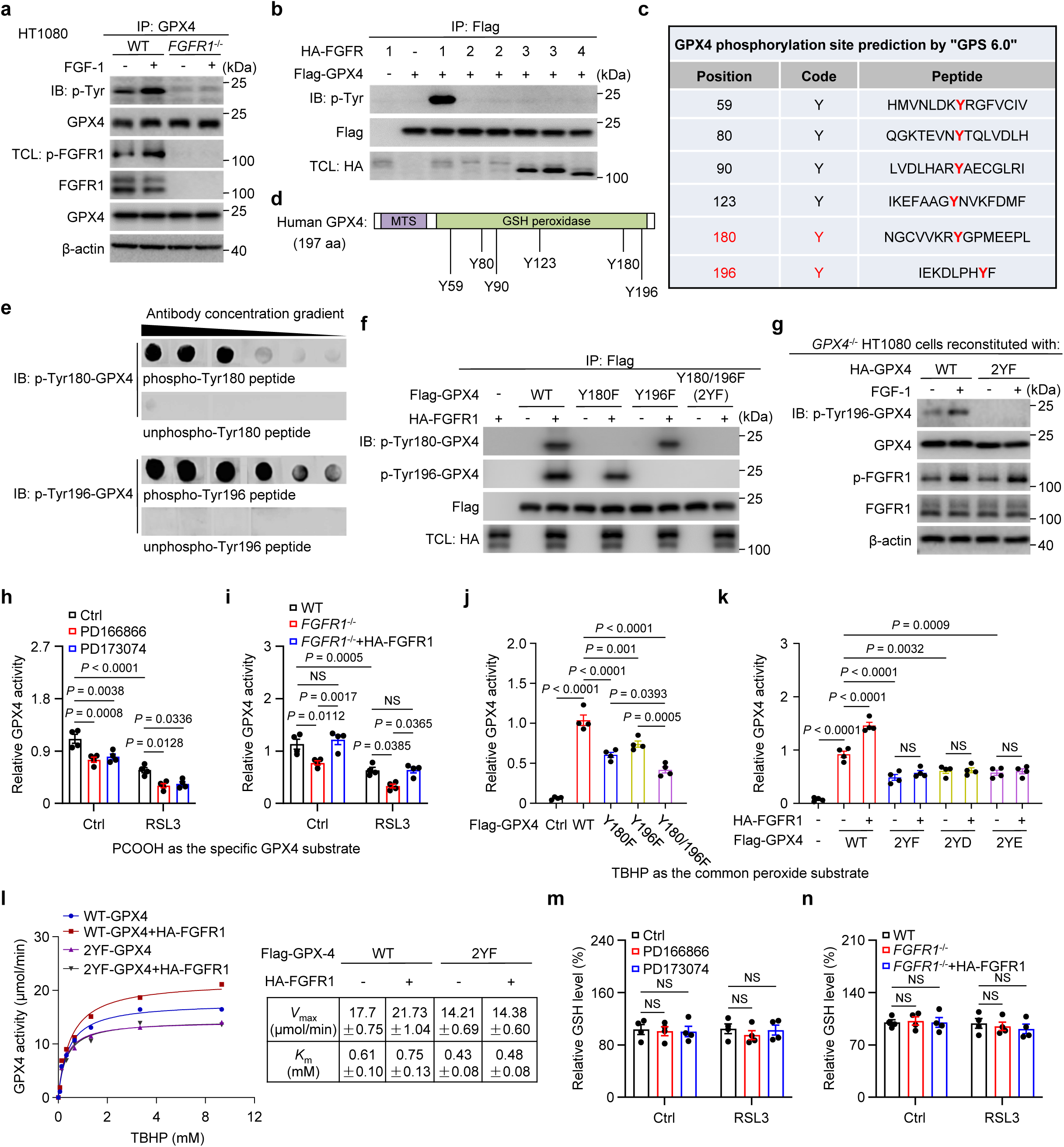
Effects of FGFR1-mediated phosphorylation on GPX4 activity and ferroptosis. **a,** Increased FGFR1-dependent tyrosine phosphorylation of GPX4 upon FGF-1 treatment. WT and *FGFR1*^−/−^ HT1080 cells were maintained in serum-free medium for 4 h, followed by stimulation with or without FGF-1 for 0.5 h. GPX4 was immunoprecipitated and analyzed by immunoblotting. **b,** FGFR1 specifically phosphorylates GPX4 among the examined FGFR family members. **c,d,** Prediction of potential tyrosine phosphorylation sites on human GPX4 by the GPS 6.0 program. MTS, mitochondrial targeting sequence. **e,** The anti-p-Tyr180 or p-Tyr196 antibodies specifically recognized phosphorylated GPX4. Keyhole limpet hemocyanin (KLH)-tagged oligo-peptides containing phosphorylated Tyr180 or Tyr196 residues were used to immunize rabbits, followed by purification of anti-GPX4-phospho-Tyr180 or –Tyr196 antibodies (designated as p-Tyr180-GPX4 or p-Tyr196-GPX4) with CNBr beads. The specificity of p-Tyr180-GPX4 or p-Tyr196-GPX4 was determined by detecting KLH-tagged peptides with or without Tyr180 or Tyr196 phosphorylated by ELISA. **f,** The specificity of our homo-made p-Tyr-180-GPX4 and p-Tyr-196-GPX4 antibodies were confirmed by immunoblot with this antibody. The indicated expression plasmids were transfected into HEK293T cells with or without HA-FGFR1. **g,** The 2YF-GPX4 mutant fails to be phosphorylated in FGF-1-stimulated in HT1080 cells. *GPX4*^−/−^HT1080 cells stably expressing HA-tagged WT-GPX4 or 2YF-GPX4 were treated with or without FGF-1. **h,i,** GPX4 activity in HT1080 cells treated with or without FGFR1 inhibitors (**h**), and in WT or *FGFR1*^−/−^ HT1080 cells reconstituted with vector or HA-FGFR1 (**i**). 1-palmitoyl-2-hydroperoxyoctadecadienoyl-sn-glycero-3-phosphocholine (PCOOH) was used as the specific substrate for GPX4 activity. **j,** GPX4 activity assay in HEK293T cells transfected with the indicated vectors. HEK293T cells were transfected with empty vector (Ctrl), Flag-tagged WT-GPX4, Y180F-GPX4, Y196F-GPX4, or Y180/196F (2YF) double-mutant GPX4. Flag-GPX4 was immunoprecipitated, and its enzymatic activity was measured using tert-butyl hydroperoxide (TBHP) as the substrate. **k,** FGFR1-mediated phosphorylation enhances GPX4 activity. Flag-tagged WT-GPX4, 2YF-GPX4, 2YD-GPX4, or 2YE-GPX4 was co-transfected with or without HA-FGFR1 into HEK293T cells. Flag-GPX4 was immunoprecipitated, and its enzymatic activity was measured with TBHP as the substrate. **l,** Enzymatic activities of WT or 2YF form of GPX4 co-expressed with or without HA-tagged FGFR1 in HEK293T cells. Michaelis constants (Km) were determined from activities generated using at least 8 concentrations of TBHP substrates as indicated (*n* = 3 assays). **m,n,** Relative glutathione (GSH) levels in primary mouse cardiomyocyte treated with or without FGFR1 inhibitors (**m**), and WT or *FGFR1*^−/−^ HT1080 cells reconstituted with vector or HA-FGFR1 (**n**). *n* = 3-4 biologically independent experiments in (**h-n**). Data are mean ± s.e.m.; ordinary two-way ANOVA, followed by Tukey in (**h, i**); ordinary one-way ANOVA, followed by Tukey in (**j, k**);ordinary two-way ANOVA, followed by Sidak in (**l-n**); \**p* < 0.05, \*\**p* < 0.01, \*\*\**p* < 0.001, \*\*\*\**p* < 0.0001, NS, not significant; Gel source data blots can be found in Supplementary Fig. 1.

**Extended Data Fig. 10.**
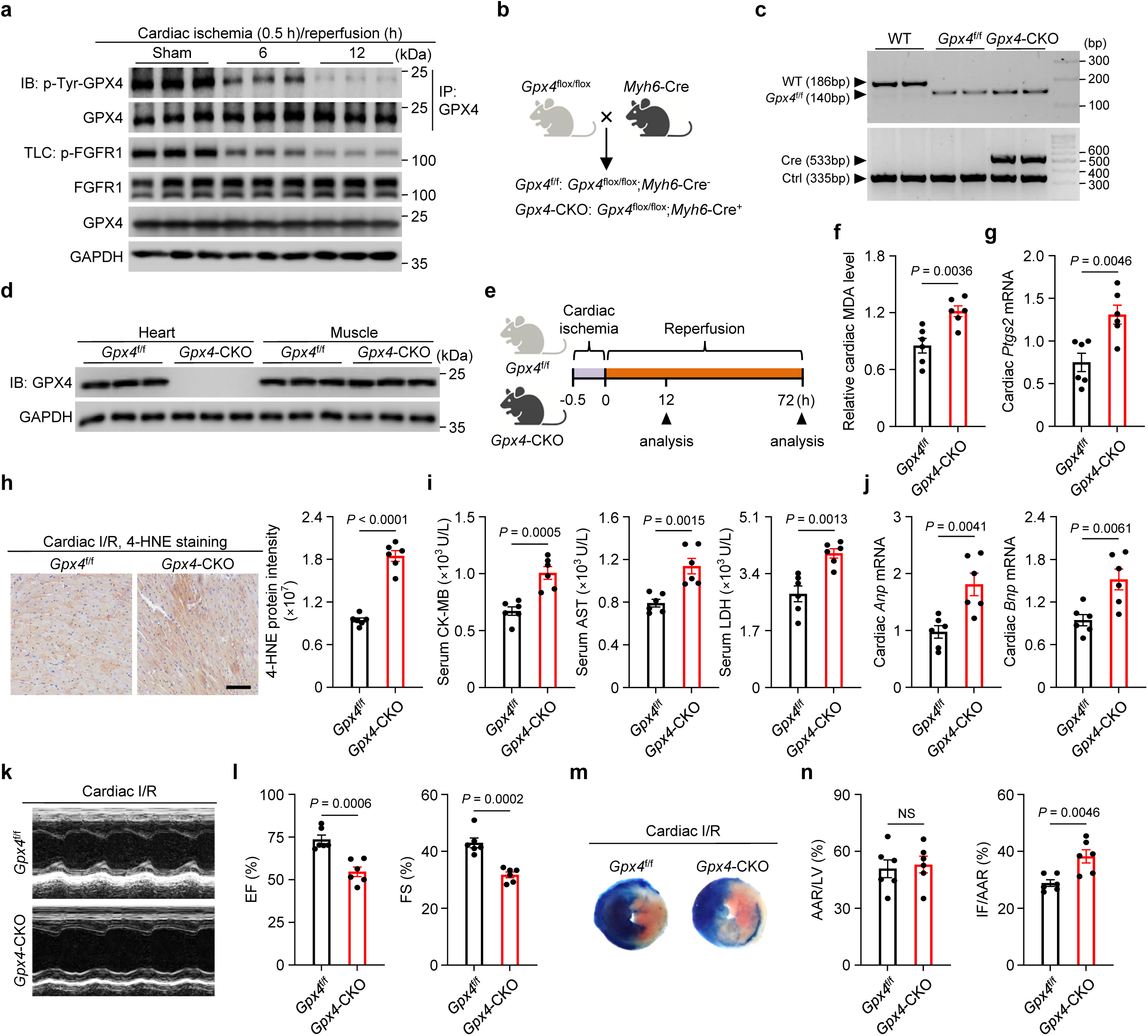
*Gpx4* deficiency aggravated cardiac I/R-induced myocardial injury by promoting ferroptosis. **a,** Immunoblot analysis of total cardiac lysates from C57BL/6J mice collected at the indicated time after cardiac I/R treatment. **b,** Schematic representation of cardiac-specific *Gpx4* knockout (*Gpx4*-CKO) mice. **c,d,** PCR genotyping (**c**) and immunoblot analysis (**d**) confirming successful cardiac-specific *Gpx4* knockout in mice. **e,** Diagram depicting the progression of cardiac I/R-induced myocardial injury in *Gpx4*^f/f^ and *Gpx4*-CKO mice. **f,** Relative MDA levels in *Gpx4*^f/f^ and *Gpx4*-CKO mice after cardiac I/R injury. **g,** Relative cardiac *Ptgs2* mRNA in the indicated mice. **h,** Representative images and quantification of 4-HNE–stained cardiac sections from the indicated mice. Scale bar, 100 μm. **i,** Serum CK-MB, AST, and LDH levels in *Gpx4*^f/f^ and *Gpx4*-CKO mice subjected to cardiac I/R injury. **j,** Cardiac *Anp* and *Bnp* mRNA in the indicated mice subjected to cardiac I/R. **k,l,** Representative echocardiography images (**k**) and statistical analysis (**l**) of EF and FS after cardiac I/R injury. **m,n,** Representative images (**m**) and quantitative data (**n**) for relative area at risk (AAR) and infarct size (IF) in heart sections from *Gpx4*^f/f^ and *Gpx4*-CKO mice subjected to cardiac I/R injury. *n* = 6 mice/group in (**f-j, l, n**). Data are mean ± s.e.m.; Two-tailed unpaired Student *t* test (**f, g, i, j, l, n,** right graph of **h**); \*\**p* < 0.01, \*\*\**p* < 0.001, \*\*\*\**p* < 0.0001, NS, not significant; Gel source data blots can be found in Supplementary Fig. 1.

**Extended Data Fig. 11.**
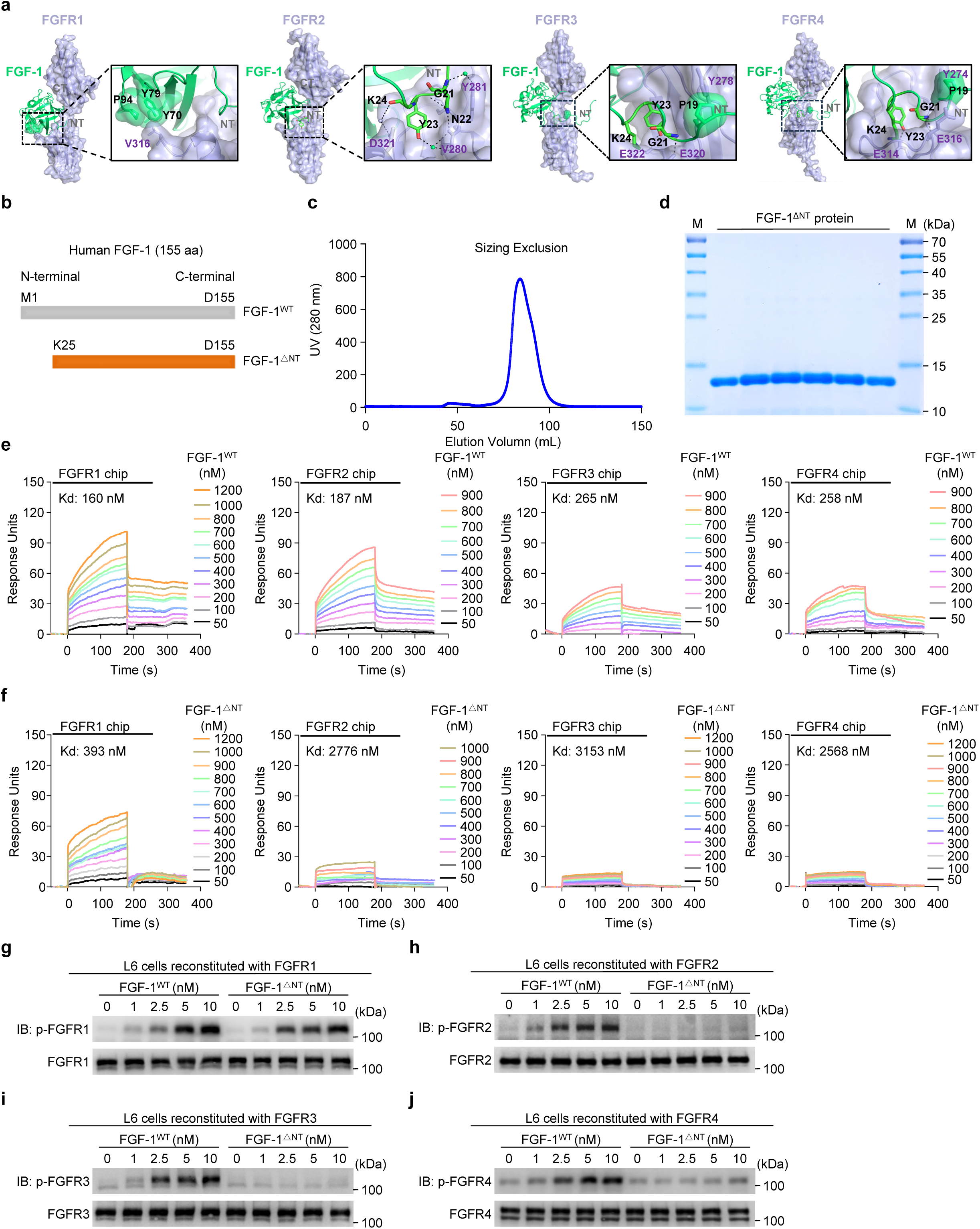
Generation of a genetically engineered recombinant human FGF-1 mutant with selectivity for FGFR1 binding and activation. **a,** Cartoon representation of the FGF-1-FGFR complex crystal structures: FGF-1-FGFR1 (PDB: 3OJV), FGF-1-FGFR2 (PDB: 1DJS), or FGF-1-FGFR3 (PDB: 1RY7). The FGF-1-FGFR4 complex crystal structure was predicted using AlphaFold 3 based on the crystal structure of the FGF-1 (PDB: 1RY7) and FGFR4 (PDB: 7YSW). **b,** Schematic illustration of human wild-type FGF-1 (FGF-1^WT^, Met1-Asp155) and the N-terminally truncated FGF-1 variant (FGF-1^ΔNT^, Lys25-Asp155). NT, N-terminal. **c,** Purification of FGF-1^ΔNT^ by gel filtration. **d,** Analysis of the fractions contained in the major peak shown in (**c**) by coomassie brilliant blue staining. **e,f,** Representative SPR sensorgrams showing the binding of FGF-1^WT^ (**e**) and FGF-1^ΔNT^ (**f**) to the ligand-binding domains of FGFR1, FGFR2, FGFR3 and FGFR4. Equilibrium dissociation constants (Kd values) were derived from saturation binding curves. **g-j,** Immunoblots showing dose-dependent activation of FGFRs by FGF-1^WT^ and FGF-1^ΔNT^ in the L6 cells expressing FGFR1 (**g**), FGFR2 (**h**), FGFR3 (**i**), or FGFR4 (**j**).Gel source data blots can be found in Supplementary Fig. 1.

**Extended Data Fig. 12.**
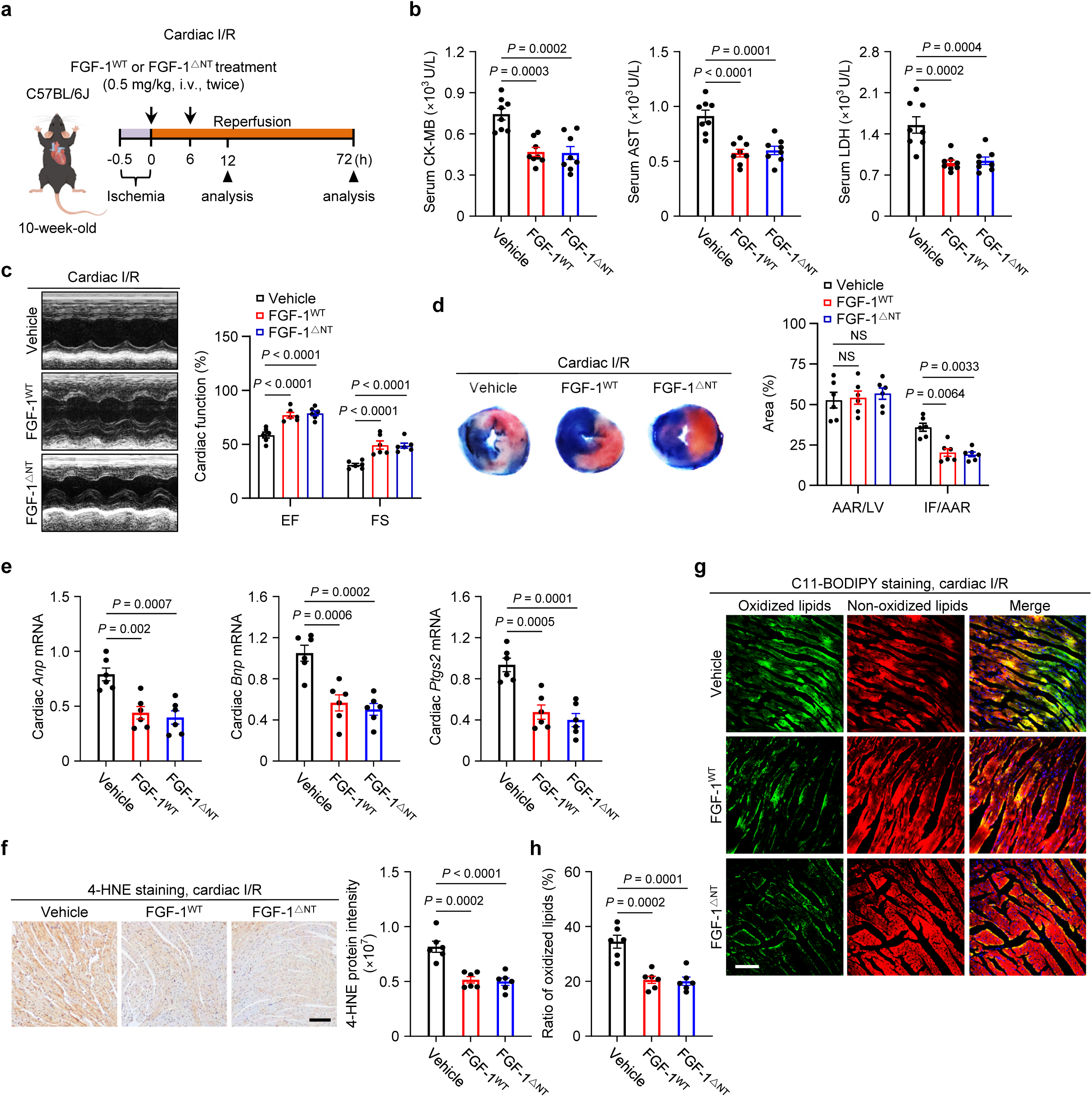
The selective FGFR1 agonist FGF-1^△NT^ attenuates cardiac I/R injury through the inhibition of ferroptosis. **a,** Schematic of cardiac I/R injury model treated with FGF-1. C57BL/6 mice were subjected to cardiac I/R, followed by i.v. administration of FGF-1^WT^ or FGF-1^△NT^ (0.5 mg/kg) at the onset of reperfusion and 6 h post-reperfusion. **b,** Serum CK-MB, AST and LDH levels in the indicated mice subjected to cardiac I/R injury. *n* = 8 mice/group. **c,** Representative echocardiography images and statistical analysis of EF and FS in the indicated mice after cardiac I/R injury. **d,** Representative images (left) and quantitative data (right) for relative AAR and IF in heart sections obtained from the indicated mice subjected to cardiac I/R injury. **e,** Relative mRNA levels of cardiac *Anp*, *Bnp*, and *Ptgs2* in the indicated mice. **f,** Representative images and quantification of 4-HNE–stained cardiac sections from the indicated mice. Scale bar, 100 μm. **g,h,** Representative images (**g**) and quantification (**h**) of C11-BODIPY staining in heart sections from the indicated groups. Scale bar, 100 μm. (**c-f, h**) *n* = 6 mice/group. Data are mean ± s.e.m.; ordinary one-way ANOVA, followed by Tukey in (**b, e,** right graph of **f**); ordinary two-way ANOVA, followed by Sidak in (right graphs of **c** and **d**); \*\**p* < 0.01, \*\*\**p* < 0.001, \*\*\*\**p* < 0.0001, NS, not significant.

**Extended Data Fig. 13.**
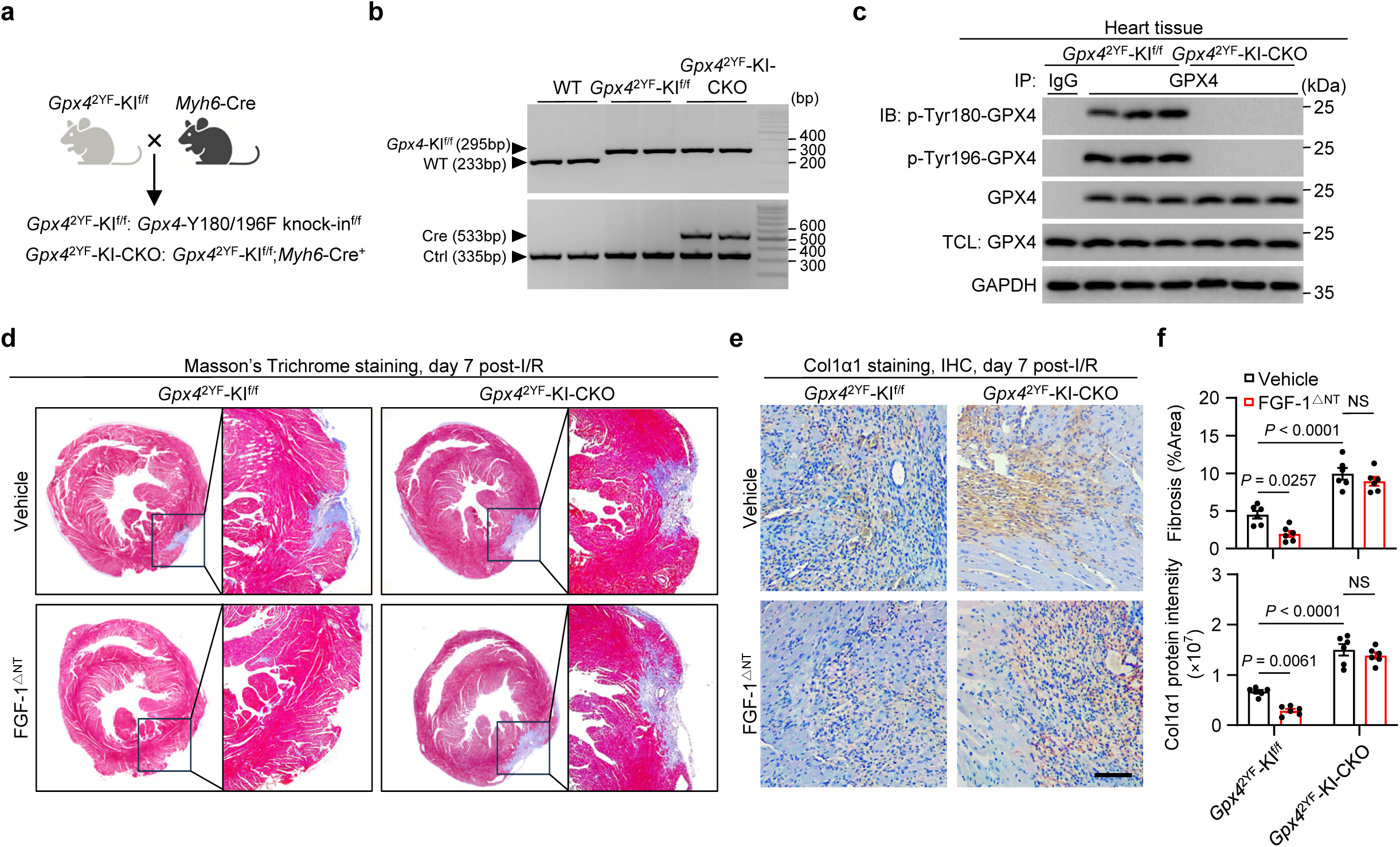
Validation of cardiomyocyte-specific *Gpx4*-Y180/196F knock-in mice. **a,** Schematic of the strategy used to generate cardiac-specific *Gpx4*-Y180/196F knock-in (*Gpx4*^2YF^-KI-CKO) mice. **b,c,** PCR genotyping (**b**) and immunoblot analysis (**c**) confirming successful cardiac-specific *Gpx4*-Y180/196F knock-in in mice. **d-f,** Representative images (**d, e**) and quantification (**f**) of Masson’s trichrome (**d**) or Col1α1-stained (**e**) heart sections from indicated mice at day 7 post-I/R. Scale bar, 100 μm. (**f**) *n* = 6 mice/group. Data are mean ± s.e.m.; ordinary two-way ANOVA, followed by Tukey in (**f**); \**p* < 0.05, \*\**p* < 0.01, \*\*\*\**p* < 0.0001, NS, not significant; Gel source data blots can be found in Supplementary Fig. 1.

**Extended Data Fig. 14.**
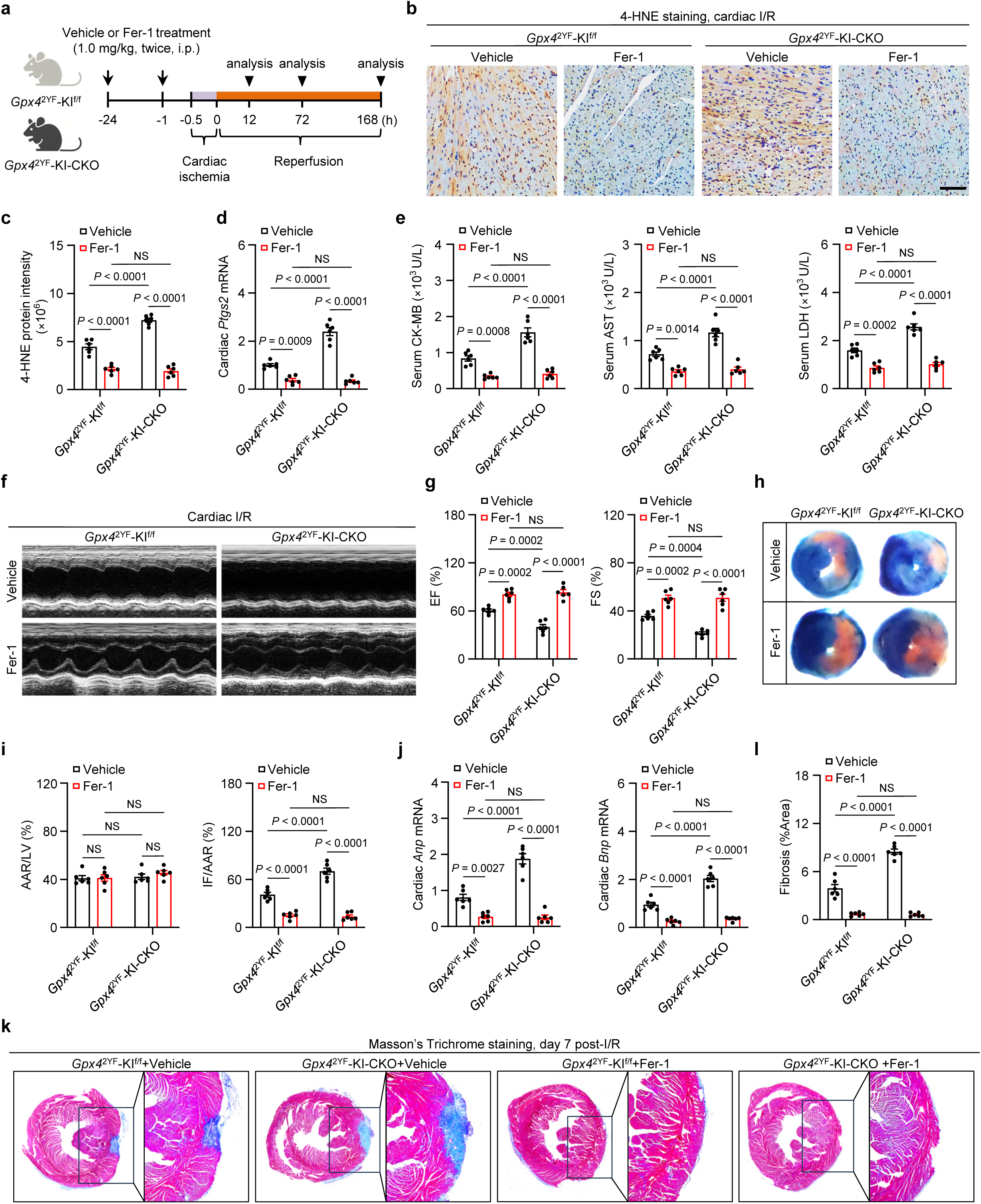
Pharmacological inhibition of ferroptosis attenuates cardiac I/R. **a,** Schematic of the ferroptosis inhibitor Fer-1 treatment regimen for *Gpx4*^2YF^*-*KI^f/f^ and *Gpx4*^2YF^-KI-CKO mice with cardiac I/R injury. Mice received Fer-1 or vehicle at the 24 h and 0.5 h pre-ischemia, followed by cardiac I/R. **b,c,** Representative images (**b**) and quantification (**c**) of 4-HNE–stained cardiac sections from the indicated mice. Scale bar, 100 μm. **d,** Cardiac *Ptgs2* mRNA in the indicated mice. **e,** Serum CK-MB, AST and LDH levels in the indicated mice. **f,g,** Representative echocardiography images (**f**) and statistical analysis of EF and FS (**g**) in the indicated mice after cardiac I/R injury. **h,i,** Representative images (**h**) and quantitative data (**i**) for relative AAR and IF in heart sections obtained from the indicated mice. **j,** Relative mRNA levels of cardiac *Anp* and *Bnp* in the indicated mice. **k,l** Representative images (**k**) and quantification (**l**) of Masson’s trichrome staining in heart sections from the indicated mice at day 7 post-I/R. (**c-e, g, i, j, l**) *n* = 6 mice/group. Data are mean ± s.e.m.; ordinary two-way ANOVA, followed by Tukey in (**c-e, g, i, j, l**); \**p* < 0.05, \*\**p* < 0.01, \*\*\**p* < 0.001, \*\*\*\**p* < 0.0001, NS, not significant.

