## Supplemental Table 1-4 for "A GPX4 phosphorylation switch by FGFR1 guards against ferroptosis"

**Supplementary Table S1. Primer sequences for genotype identification of mice**

| Genotype | Sequence (5'-3') |
|---|---|
| *Fgfr1*$^{flox/flox}$ | Forward: TTGTAGAGTGGAATTGGTTGCCAG |
| | Reverse: GTATGTCCCATGTGTATGCAGGA |
| *Gpx4*$^{flox/flox}$ | Forward-WT: CTGCAACAGCTCCGAGTTC |
| | Forward-MUT: CCAGTAAGCAGTGGGTTCTC |
| | Reverse: CGGTGCCAAAGAAAGAAAGT |
| *Gpx4*-Y180/196F knock-in$^{flox/flox}$ | Forward: GCAAGTCTGTGTCATGCATGCAT |
| | Reverse: GGCAAGTGGATATGAGAGTTCTAAGC |
| *Myh6*-Cre | Forward: GAAATGACAGACAGATCCCTCCTATC |
| | Reverse: CGACGATGAAGCATGTTTAGCTG |
| Internal control for *Myh6*-Cre | Forward: CATGCCAATGGTTCACTCTAAGGT |
| | Reverse: TCTCTATGTCCCAAAGTGCAGACAC |

**Supplementary Table S2. Primer sequences for identification of sgRNA**

| Gene | Sequences (5'-3') |
|---|---|
| Human *FGFR1*_sg1 | Forward: TGGGCAAGACACCTCCAG<br>Reverse: CGTACCTTGTAGCCTCCAATTC |
| Human *FGFR1*_sg2 | Forward: GGTTTCATCTGAGAAGCAAGGA<br>Reverse: AGGGAGAAAGCAGGACTCTACA |
| Human *GPX4*_sg2 | Forward: TGTTCTTCTGCGCTGACGC<br>Reverse: GCGAACTCTTTGATCTCTTCGT |
| Mouse *Fgfr1*_sg1 | Forward: GACCAGTACTCACCCAGCTTTC<br>Reverse: AAAGGCAGTGTGTCCAACAAG |
| Mouse *Fgfr1*_sg2 | Forward: AGAAGGGAGTCCTGGAGAAAAG<br>Reverse: GCTCCCAGAAGGTTGATGATA |
| Mouse *Plcγ*_sg1 | Forward: ACTCTGAGTTTGACAGCCTGGT<br>Reverse: ATTTCCTCCAGCTACACACCAT |
| Mouse *Plcγ*_sg2 | Forward: GCACCGTCATGACTTTGTTCTA<br>Reverse: GGTGTCCTATCAGGTGATCCAA |
| Mouse *Frs2*_sg1 | Forward: TATACAGGTGGATTTGGAAGGG<br>Reverse: TAGACATGAAGAGGAGGCACTG |
| Mouse *Frs2*_sg2 | Forward: CATTAATGTGGATGATGATGGG<br>Reverse: ACCTTGTCCAGTCTGACACCTT |

**Supplementary Table S3. Primer sequences for qRT-PCR**

| Gene Name | Sequence (5'-3') |
|---|---|
| Mouse-*Fgfr1* | Forward: ACTCTGCGCTGGTTGAAAAAT |
| | Reverse: GGTGGCATAGCGAACCTTGTA |
| Mouse-*Fgfr2* | Forward: CCACCTCGATGTCGTTGAAC |
| | Reverse: GCCCATCAGGCCCGTATTTAC |
| Mouse-*Fgfr3* | Forward: GCCTGCGTGCTAGTGTTCT |
| | Reverse: CCTGTACCATCCTTAGCCCAG |
| Mouse-*Fgfr4* | Forward: TCCATGACCGTCGTACACAAT |
| | Reverse: ATTTGACAGTATTCCCGGCAG |
| Mouse- *Ptgs2* | Forward: TGCACTATGGTTACAAAAGCTGG |
| | Reverse: TCAGGAAGCTCCTTATTTCCCTT |
| Mouse-*Anp* | Forward: GTGCGGTGTCCAACACAGAT |
| | Reverse: TCCAATCCTGTCAATCCTACCC |
| Mouse-*Bnp* | Forward: GAGGTCACTCCTATCCTCTGG |
| | Reverse: GCCATTTCCTCCGACTTTTCTC |
| Mouse-*Myh7* | Forward: AGACTGTCAACACTAAGAGGGT |
| | Reverse: TGCCCCAAAATGGATTCGGAT |
| Mouse-*Gapdh* | Forward: AATGTGTCCGTCGTGGATCT |
| | Reverse: CATCGAAGGTGGAAGAGTGG |

**Supplementary Table S4. Primer sequences for molecular cloning**

| Gene | Forward primer (5'-3') | Reverse primer (5'-3') |
|---|---|---|
| FGFR1 | ATGTGGAGCTGGAAGTGCC | TCAGCGGCGTTTGAGTCC |
| FGFR1-Y653/654F | ATCGACTTCTTTAAAAAGA CAACCAACGGCC | CTTTTTAAAGAAGTCGATG TGGTGAATGTC |
| FGFR1-Y463F | TCTGAGTTTGAGCTTCCCG AAGACCCT | AAGCTCAAACTCAGAGACC CCTGCTAG |
| FGFR1-Y583F | CTGGAATTCTGCTACAACC CCAGCCACAACC | GTAGCAGAATTCCAGCCCT GGGGGCC |
| FGFR1-Y585F | TACTGCTTCAACCCCAGCC ACAACCCAGAG | CCAGGGCTGGAATACTGCT TCAACCCC |
| FGFR1-Y730F | GAGCTGTTCATGATGATGC GGGACTGCTGGCA | CATCATGAACAGCTCGTTG GTGCAGTTACTG |
| FGFR1-Y766F | CAGGAGTTCCTGGACCTGT CCATGCCC | GTCCAGGAACTCCTGGTTG GAGGTCAAG |
| FGFR1 (aa1-357) | ATGTGGAGCTGGAAGTGCC | GGTCAACCATGCAGAGTGA T |
| FGFR1 (aa478-822) | CTGGTCTTAGGCAAACCC | TCAGCGGCGTTTGAGTCC |
| FGFR1 (aa1-596) | ATGTGGAGCTGGAAGTGCC | GGAGAGCTGCTCCTCTGG |
| FGFR1 (aa597-767) | TCCAAGGACCTGGTGTCC | CAGGTACTCCTGGTTGGAG |
| FGFR1 (aa597-822) | TCCAAGGACCTGGTGTCC | TCAGCGGCGTTTGAGTCC |
| FGFR1 (aa1-767) | ATGTGGAGCTGGAAGTGCC | CAGGTACTCCTGGTTGGAG |
| FGFR2 | ATGGTCAGCTGGGGTCGTT | TCATGTTTTAACACTGCCG TT |
| FGFR3 | ATGGGCGCCCCTGCCTG | TCACGTCCGCGAGCCCC |
| FGFR4 | ATGCGGCTGCTGCTGGC | TCATGTCTGCACCCCAGAC |
| GPX4 (aa1-197) | ATGAGCCTCGGCCGCCTT | CTAGAAATAGTGGGGCAGG T |
| GPX4 (aa1-267) | ATGAGCCTCGGCCGCCTT | CTAATTTGTCTGTTTATTC |
| GPX4-Y180F | AAGCGCTTCGGACCCATGG AGGAGCC | GGGTCCGAAGCGCTTCACC ACGCAGC |
| GPX4-Y196F | CCCCACTTTTTCTAGCTCG AGCCATGGAA | CTAGAAAAAGTGGGGCAGG TCCTTCTC |

| | | |
|---|---|---|
| GPX4-Y180D | AAGCGCGACGGACCCATGGAGGAGCC | GGGTCCGTCGCGCTTCACCACGCAGC |
| GPX4-Y196D | CCCCACGACTTCTAGCTCGAGCCATGGAA | CTAGAGTCAGTGGGGCAGGTCCTTCTC |
| GPX4-Y180E | AAGCGCGAAGGACCCATGGAGGAGCC | GGGTCCTTCGCGCTTCACCACGCAGC |
| GPX4-Y196E | CCCCACGAATTCTAGCTCGAGCCATGGAA | CTAGATTCAGTGGGGCAGGTCCTTCTC |
| FGF-$1^{WT}$ (aa1-155) | ATGGCTGAAGGGGAAATCAC | TTAATCAGAAGAGACTGGCA |
| FGF-$1^{\Delta NT}$ (aa25-155) | AAGCCCAAACTCCTCTACTG | TTAATCAGAAGAGACTGGCA |
| FGFR1 (aa142-365) | ATGGATAACACCAAACCAAAC | TCACCTCTCTTCCAGGGCT |
| FGFR2 (aa149-368) | ATGAACAACAAGAGAGCACC | TCACTCCTTTTCTCTTCCAG |
| FGFR3 (aa147-365) | ATGAAGGGGCCCCTGTGTC | TCACTCCACCAGCTCCTC |
| FGFR4 (aa144-355) | ATGCAGCAAGCACCCTACTG | TCAGTCCTCCTCTGGCAGC |
| **CC155-FGFR1**： | | |
| Signal peptide | ATGTGGAGCTGGAAGTGCC | / |
| Signal peptide-CC155-FGFR1 | GTTCTTCTGCTTGTCAGCGGTGCAGAGTGTGGC | ACACTCTGCACCGCTGACAAGCAGAAGAACGGCAT |
| FGFR1 | / | TCAGCGGCGTTTGAGTCC |
| **VN155-GPX4**： | | |
| GPX4 | ATGAGCCTCGGCCGCCTT | / |
| VN155-GPX4 | GGATCCACTGAATTCGAAATAGTGGGGCAGGTCC | CTGCCCCACTATTTCGAATTCAGTGGATCCGGAGG |
| VN155 | / | GGCGGTGAGATAGACGTTG |
